# From Body Hulls to Musculoskeletal Models: Personalized Inertial Parameter Estimation

**DOI:** 10.1101/2025.07.08.663673

**Authors:** Markus Gambietz, Putri Qistina Azam, Philipp Amon, Iris Wechsler, Eva Maria Hille, Timo Menzel, Tabea Ott, Mario Botsch, Matthias Braun, Jörg Miehling, Katie L. McMahon, Anne D. Koelewijn

## Abstract

Every human body is different, however, current movement analysis does not reflect that, as it heavily relies on generic musculoskeletal models. Usually, these models are scaled to match the participants’ body segment lengths and body weight, but not taking individual body shape into account. This can lead to errors in the estimation of joint forces and torques, which are important to accurately estimate musculoskeletal variables. Thus, we developed a method to estimate body segment inertial parameters based on body hulls acquired via smartphone pictures. From the body hull, we infer the skeletal shape and pose, and then estimate the distribution of bone, lean, and fatty tissues. We then segment the body hull and assign each tissue type a density, which is used to calculate the body segment inertial parameters. To allow for the use of our method with existing data, we also introduce two new generic musculoskeletal models, which are based on the average standing body shapes. Validation using MRI-derived ground-truth models shows that our method creates participant-specific musculoskeletal models that are closer to the MRI-derived ground truth than scaled generic models. Additionally, we performed lab-based gait experiments to evaluate the effect of our method on residual forces and joint moments, where we found that our method leads to a reduction of residual forces of up to 14.9 % and a reduction of metabolic cost of up to 12.8 % when compared to generic musculoskeletal models. Our new generic models show similar joint moment outcomes, but less reduction of residual forces than the personalized models.

## Introduction

Quantitative human movement analysis is important for many applications, such as rehabilitation, sports science, and ergonomics [1, 2]. Often, muscle and joint forces are of interest, which can be estimated using musculoskeletal (MSK) models. To avoid introducing errors, numerical parameters in MSK models need to be correct, which includes body segment inertial parameters (BSIPs) as well as joint and muscle geometry. Parameters will vary between individuals thus personalization of MSK model parameters is important.

It is not trivial to verify the *correctness* of MSK model parameters, as they are often not directly measurable or require medical imaging, which is expensive and time-consuming. However, when MSK model parameters are *incorrect*, errors in kinematics and dynamics can be expected [3]. Then, a motion reconstructed via inverse kinematics (IK) and inverse dynamics (ID) will result in residual forces, which are the difference between the measured external forces and those estimated by the MSK model. As a measure of validity, thresholds were introduced, below which residual forces are considered acceptable [4]. To comply with these thresholds, one can use residual reduction methods that optimize the MSK model parameters to minimize residual forces, such as OpenSim’s Residual Reduction Algorithm [5] or AddBiomechanics’ dynamics fit [6]. However, it is not guaranteed that the resulting parameters are actually reflecting the true inertial parameters of the participant. In residual reduction methods, many local minima can be expected, as the optimization problem is non-convex, and there are hundreds of parameters to optimize. Thus, whether using residual reduction methods or not, a good initial MSK model personalization is essential.

Current MSK model personalization is based on scaling a generic model to match the participants’ body segment lengths and full body weight, either based on static [5] or dynamic [6] marker data. This procedure entails two assumptions: first, that the generic model is a general representation of a diverse set of human bodies. Historically, BSIPs were derived from cadaveric studies with only a few participants, often with a narrow range of body shapes and sizes and a bias towards geriatric, white, male individuals [7, 8]. Current MSK models (e.g., [5, 9, 10]) draw their BSIP data from stereophotometry and gamma-ray scanning studies on young, male, caucasian populations [11–13]. Thus, they might be insufficent for the widespread use that we are seeing today, fitting van der Kruk’s findings that gender biases are prevalent in biomechanics, with males taken as the default [14]. Secondly, the commonly used length-based scaling approach ignores individual differences in body composition and segmental mass distribution because it relies solely on distances between anatomical markers rather than actual body shape or volume. Together, these limitations challenge the accuracy of personalized models and the ability to gain subject-specific biomechanical insights.

To improve the accuracy of BSIPs, we aim to take individual body shape into account. A person’s body shape, or hull, correlates with the occurence of tissue types, and therefore densities. For example, body hulls were used to estimate body fat percentage [15], skeletal shape [16, 17], or tissue distribution [18]. Body hull recording has recently become easily accessible through smartphone applications, where scanning can be completed in a few minutes without any additional equipment [19–21]. Furthermore, personalizing a human body model by shape aligns with the call from ethical and legal scholars, to consider the importance of bodily variation and aspects of gender, sex, ethnicity, or skin color [22–25].

To this end, we propose a **s**hape-based **i**nertial **p**arameter **p**ersonalization (SIPP) method, which uses an individual’s body hull to estimate their BSIPs. We acquire the body hull using a smartphone app [21], though alternative acquisition technologies, such as 3D scanning, are equally feasible. Then, we fit the Skinned Multi-Person Linear (SMPL) model [26] to the body hull. We then apply SKEL, a parametric model that infers skeletal shape and joint positions from body surface on the SMPL fit [17]. We obtain the distribution of lean and fatty tissues through the Human Implicit Tissues (HIT) model [18], which is trained on MRI data to estimate tissue occurence based on body shapes. We segment the body hull based on SKEL’s pose output and assign each tissue a density, which is then used to calculate the BSIPs. The resulting BSIPs are then inserted into a traditionally scaled MSK model (see Fig. 1). To validate our method, we aquired MRI, body hull and gait data for a small cohort, where we used the MRI to generate a ground-truth MSK model. As part of the dataset, we also include a benchmark that evaluates the quality of personalization methods when compared to MRI.

**Fig 1.**
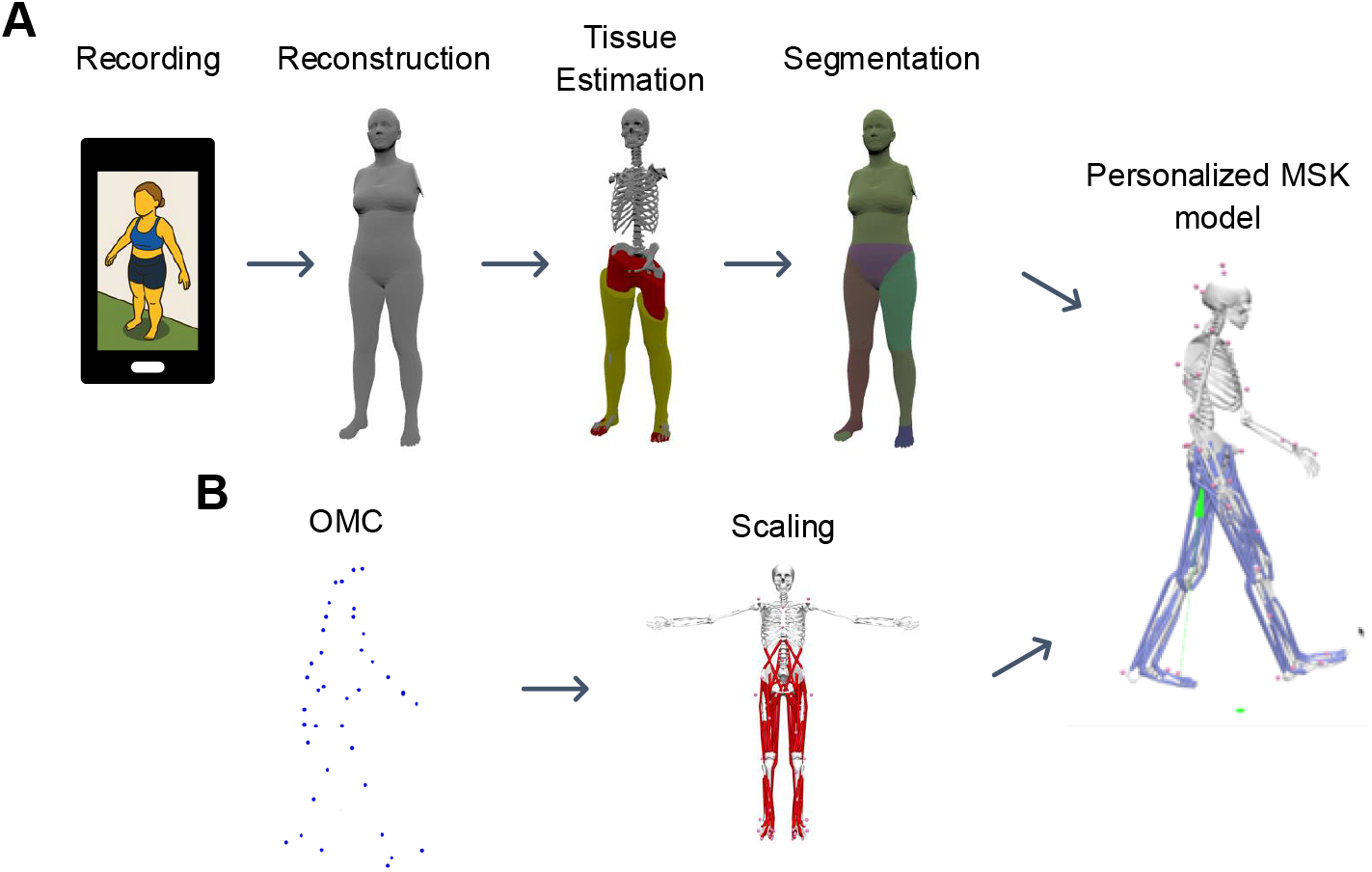
Workflow for creating personalized musculoskeletal models using smartphone-based body hulls. **A** shows the automated SIPP pipeline, which takes a body hull or smartphone pictures as input and outputs participant-specific BSIPs. **B** displays the current optical motion capture data workflow that scales a generic model to the participant’s body segment lengths, for which we use AddBiomechanics [6]. We then combine the BSIPs from SIPP with the scaled model to create a personalized MSK model. When using smartphone pictures, arms were removed from the body hull as they are affected by sway during recording.

Furthermore, we introduce two new generic MSK models, *SIPP-generic-female* and *SIPP-generic-male*. These models are variants of the Rajagopal2016 model [10], which were scaled to the average SMPL shape for each sex. We then ran SIPP on both these models to obtain their reparametrized BSIPs. *SIPP-generic-female* represents a female person with a body height of 1.66 m and a bodymass of 72 kg, whereas *SIPP-generic-male* represents a male person with a body height of 1.79 m and a bodymass of 86 kg.

In summary, our contributions are as follows:

- MRIgait dataset: A benchmark dataset containing MRI labels, body hulls reconstructed from smartphone and a commercial body scanner (scaneca.com), gait at various speeds and inclines, and MRI-personalized MSK models.
- SIPP: A shape-based inertial parameter personalization method, which we use to generate participant-specific MSK models from smartphone pictures. SIPP is meant to be used in any biomechanical assessment if body hull recordings are feasible.
- *SIPP-generic-female* and *SIPP-generic-male*: Two generic musculoskeletal model parameterizations that are based on average standing body shapes. These models are meant to be used with existing gait data or when no matching body hulls can be recorded.

We found that both SIPP and SIPP-generic reduced residuals, and that their joint torque estimates were more similar to MRI-personalized models than to scaled MSK models. Remarkably, using SIPP-parametrized models reduces metabolic cost estimates by up to 12.8 %. Furthermore, we show that residual reduction methods do not necessarily result in BSIPs that are closer to the MRI, and that SIPP in combination with residual reduction produced the lowest residuals.

## Materials and methods

The study was approved by the ethics committee of the Friedrich-Alexander-Universität Erlangen-Nürnberg (Nr. 20-471 1-B).

### Data Acquisition

We recruited 6 participants (age (27.2 *±*1.5) years, 3F - height (1.60 *±*0.09) m, mass (60.6 *±* 11.5) kg; 3M - height (1.78 *±* 0.01) m, mass (83.4 *±* 7.2) kg) to acquire gait data, smartphone-based body hulls and MRI scans. Participants were recruited from the University of Erlangen-Nürnberg (Germany) from June 12 to 30, 2024 and provided written informed consent prior to participation. All procedures were carried out in accordance with the Declaration of Helsinki. Participants were excluded if they had any known musculoskeletal or neurological disorders, implanted metallic devices, were underage, or if they were pregnant. All participants were free of any injuries at the time of the study. Due to configuration of scanner and coils, we excluded participants taller than 1.8 m.

### Body Hull

We acquired full-body scans of participants in a standardized A-pose with shoulderwide stance using a preliminary version of the ‘Avatars for the Masses’ smartphone app [21], ensuring good lighting and minimal background interference. To eliminate clothing artifacts, participants were asked to wear minimal clothing that is as tight-fitting as possible, but only as far as they felt comfortable. The app requires circling the participant while pictures are taken automatically. The participant should be in frame completely during the capture process. As we are especially interested in the lower limbs, we circled each participant a second time, with the legs in the center of the frame. If a participant moved noticeably, we repeated the recording. Each scan typically took less than 5 minutes.

### MRI

To create reference MSK models, we acquired MRI scans of the participants using a 3T MRI scanner (MAGNETOM Prisma, Siemens Healthineers, Erlangen, Germany). The scans were acquired using a T1-weighted 3D FLASH sequence with the following parameters: TR = 50 ms, TE = 1.33 ms, flip angle = 16°, voxel size = 0.8 mm x 0.8 mm x 2.0 mm, matrix size = 640 x 640 x 160, and a field of view of 500 mm. Depending on the height of the participant, we acquired the full-body scan in 4 or 5 segments, which overlapped by 0.1 m. The scan time was approximately 90 min. Participants were scanned in a supine position with their arms resting loosely besides the body. By that, the arms and shoulders were partially out of the field of vision and we discarded them from the analysis. We opted against recording the arms, as that would almost double the scan time. After the MRI scan, we also recorded body hulls using a commercial body scanner (SCANECA, Scaneca GmbH, Berlin, Germany) to obtain the standing shape of the participants. We did not use the smartphone-based body hulls for the MRI validation, as eventual processing inaccuracies or artifacts should not affect the reference MSK models. All MRI scans were taken within 1 day of the gait and body hull recordings.

### Gait

To study the effects of personalizing BSIPs on human movement dynamics, gait data was recorded using a treadmill with embedded force plates (Motek M-Gait, Motek Medical B.V., Houten, Netherlands). We recorded 3D kinematics using Vicon’s Plug-in Gait markerset and a 12-camera motion capture system (Vicon, Vicon Motion Systems Ltd., Oxford, UK) at 100 Hz and ground reaction forces at 1000 Hz. Each participant walked on the treadmill for 5 min at 2 different speeds (0.8 ms^*−*1^, 1.3 ms^*−*1^) and 3 different inclines (0°, *−*4.57°, 4.57°). Due to problems with the equipment, a total of 5 trials were excluded (see S1 Appendix for more details), leaving us with 25 trials in total. Kinematics and scaled Rajagopal2016 MSK models were obtained using AddBiomechanics’ kinematics fit [6, 10].

### Shape-based Inertial Parameter Personalization

We reconstructed the body hull from smartphone images using the method of Wenninger et al. [19]. We found that the reconstruction of the arms would often be more affected by sway than the rest of the body, and that hair would lead to inaccurate head volumes. Therefore, we created an initial SMPL fit [26, 27], from which we identified the vertices in the body scan that correspond to the head and arms based on the closest vertex from the SMPL fit by euclidean distance. We then removed these vertices from the body hull. This step is needed to ensure that the SMPL shape parameters are not biased by eventual inaccurate arm reconstructions. We then fit the SMPL model to the pruned hull, while keeping landmarks at the wrists to retain the correct pose.

Based on the SMPL fit, we fit SKEL [17] and run HIT [18]. SKEL is a variation of SMPL that comes with biomechanically accurate joint positions and uses similar conventions as the Rajagopal2016 model [10]. Additionally, SKEL infers the skeletal shape. HIT is a method to estimate the distribution of lean, fatty, and bone tissues in the body hull, which is based on the SMPL shape parameters. Based on their outputs, we assign densities according to Table 1 for bone, muscle, lung, and fat tissues. HIT and SKEL both estimate skeletal shape, but we only use SKEL’s skeletal shape, as it is more detailed. HIT’s bony tissues are therefore merged with its lean tissue. Unoccupied space in the torso is considered to be the lung. In the torso, lean tissue includes the organs, therefore, we assign a lower density here, which is based on the average organ density.

**Table 1.** Tissue densities that were assigned to the smartphone-based body hull and MRI scan. Densities and standard deviations are based on [28], however, we could only use the mean values. For the organs, which make up the majority of the lean tissue in the torso, we uniformly assigned a density of 1050 kg m^*−*3^, corresponding to the average organ density [28]. In other parts of the body, we assigned the lean tissue the same density as muscles.

| Tissue Type | Density [$\text{kg m}^{-3}$] |
| --- | --- |
| SIPP |  |
| Lean Tissue (torso) | 1050 |
| Lean Tissue (others) | $1090 \pm 55$ |
| Adipose Tissue | $911 \pm 51$ |
| Bone | $1178 \pm 197$ |
| Lung | $394 \pm 159$ |
| MRI Scans |  |
| Organs | 1050 |
| Muscle | $1090 \pm 55$ |
| Fat | $911 \pm 51$ |
| Trabecular bone | $1178 \pm 197$ |
| Cortical bone | $1908 \pm 166$ |
| Lung | $394 \pm 159$ |

We then segment all resulting meshes based on planes that we constructed from SKEL’s pose and joint centers. For most joints, we use the orientation of the neighboring segments to construct a normal that is perpendicular to the segmentation plane. For example, the segmentation normal for the *knee* joint is defined by summing the orientation vectors of the *thigh* and *shank* segments. This procedure, however, would not lead to a physiologically plausible segmentation for the hip and shoulder joints, as their parent segment orientations are not representative of the actual joint orientation. Therefore, we construct points which are used to create the segmentation plane normal vector that points towards the joint center. For the shoulder, we set that point at 75 % of the distance between SKEL’s *lumbar* and *thorax* joints, while the hip reference point is set to be the average between the *hip*, contralateral *hip*, and *lumbar* joints. All joints are segmented at their joint centers, except for the *lumbar* joint, which is more distal in the Rajagopal2016 model. We therefore set the origin of the *lumbar* segmentation plane to be at 40 % of the distance between SKEL’s *hip* and *lumbar* joints.

Segment BSIPs were estimated from the segmented meshes and their assigned densities, rotated into OpenSim’s coordinate frames and then inserted into a scaled Rajagopal2016 model. Due to processing inaccuracies, the total body mass will not match the actual mass of the participant, but deviate by a small margin. Therefore, we scaled the mass and moment of inertia to match the actual mass of the participant as measured with force plates from the treadmill. The process to estimate BSIPs from body hulls is automated and takes approximately 5 minutes per participant on a desktop computer equipped with a GPU (NVIDIA RTX 4090). We also test the sensitivity of the internal tissue estimation model by replacing HIT with the layered tissue model “InsideHumans” [15] and by skipping the internal tissue estimation step, i.e. assuming a constant density for all tissues.

To create the *SIPP-generic-female* and *SIPP-generic-male* MSK models, we start with an A-pose and neutral shape parameters for the female and male SMPL model, respectively. We then scale the Rajagopal2016 MSK model to SMPL’s height and run the same pipeline as for the smartphone-recorded body hulls. Although SIPP-generic does not account for individual body shape, it offers an alternative to current generic BSIP parameterizations, which were derived from less accurate and less diverse body hulls and assumed constant densities [11].

### Obtaining Inertial Parameters from MRI

As we are interested in extracting BSIPs from the MRI scan, we first have to assign densities to each voxel in the scan. To accomplish this, we used TotalSegmentator [29] to annotate body hull, muscle, fat, bone, and lung tissues. Each segmentation was reviewed and corrected manually, which took several hours per participant. Lastly, we thresholded the voxel intensity to categorize the bone into cortical and trabecular sections. Densities were annotated according to Table 1.

Tissue deforms when laying on the table. To compensate for this, we use the actual standing shape of the participant, which we recorded with a commercial body scanner. Here, the goal is to find a deformation field that allows us to deform the compressed MRI body shape into the actual standing shape of the participant. We obtain the deformation field through a volume interpolation between the MRI body hull and the standing body hull. To find a valid interpolation, both body hulls need to aligned and in the same pose. Therefore, we first fit a SMPL model to the standing body hull. This model is then rigged to match the outer shape of the MRI annotations. Then, the MRI body hull is obtained by deforming the vertices of the SMPL model to match the outer shape of the MRI annotations. Now we can interpolate each point in the MRI annotations by using a cubic radial basis function interpolator [30]. Interpolator hyperparameters were tuned so that the discrepancy between the deformed and the original volumes was minimal. Any remaining volume discrepancy was compensated by scaling the resulting BSIPs. From here on, we proceed as with the smartphone-recorded body hulls.

### MRIgait Benchmark

The MRI-generated MSK models were considered to be a target outcome for any personalization, whether image- or marker-based. Therefore, we introduce the MRIgait benchmark, which consists of four metrics that compare personalized to the MRI-generated MSK models. For each metric, we calculate the mean absolute deviation (MAD) and the mean relative deviation (MRD) between the personalized and the MRI-generated MSK models. The first three metrics are the per-segment deviations for mass, center of mass distance from the parent joint, and the diagonal axis components of the moment of inertia tensor. We chose to disregard off-axis components as they are set to zero in generic OpenSim models and would therefore skew the benchmark.

Furthermore, we combine all the toe, calcaneus, and talus segments into a single foot segment as its joints are locked during inverse kinematics. The remaining metric is the deviation in kinetic energy. Kinetic energy is calculated for every state (pose **q**, velocities 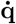) for a given participant. This metric judges functional, task-specific, personalization quality over a given dataset. During gait, for example, the lumbar joint sees very little movement, therefore the inertial parameters of the pelvis and torso are close to redundant with regard to kinetic energy. In this case, incorrect mass assignment between the pelvis and torso would not be punished. The kinetic energy metric can also be used to test BSIP generalization by using general datasets such as BioAMASS [17] or the AddBiomechanics dataset [31].

### Gait Data Processing and Comparisons

We filtered ground reaction forces and kinematics at 15 Hz (2nd order bi-directional butterworth filter) before running inverse dynamics using OpenSim. In the first step, we compared all resulting MSK models on the MRIgait benchmark. Then, we compared the same outcome variables from AddBiomechanics’ physics optimization method that are either initialized with a generic (AddBiomechanics), SIPP-personalized (AddBiomechanics+SIPP), or SIPP-generic (AddBiomechanics+SIPP-generic) MSK model. We re-ran inverse dynamics in OpenSim for all models, as numerical differentiation is treated differently between OpenSim and AddBiomechanics. Finally, we compared residual forces, joint moments, and metabolic costs between generic, SIPP-personalized, and SIPP-generic MSK models.

## Results

### MRIgait Benchmark Results

The results of the MRIgait benchmark are shownc in Table 2. SIPP and SIPP-generic models show a lower MAD for three out of four and a lower MRD for all four parameters compared to the scaled model. For SIPP, the reduction in deviation ranges from 19.0 % for inertia MAD up to 74.1 % for center of mass MRD. The SIPP-generic model shows similar results, with reductions ranging from 9.5 % for inertia MAD to 80.4 % for mass MRD. All relative deviations improved by more than 50 % when using SIPP or SIPP-generic compared to the scaled model. Through the use of AddBiomechanics’ physics optimization, the mass and energy metrics are improved compared to the scaled model, however, the center of mass and inertia metrics are hardly affected. When applying AddBiomechanics’ physics optimization to SIPP, no overall improvement of the metrics compared to SIPP alone was observed, while the SIPP-generic model showed a reduction in mass, inertial, and energy MRD after optimization with AddBiomechanics.

**Table 2.** Results of the MRIgait benchmark. The first two rows show state-of-the-art methods of scaling (scaled) and AddBiomechanics’ physics optimization (addB). The next two rows show the novel SIPP-personalized and scaled SIPP-generic (S-gen) models. In the final two rows, SIPP and SIPP-generic in combination with AddBiomechanics’ physics optimization are shown (add+SIPP, add+S-gen). The columns show the mean and standard deviation of the mean absolute deviation (MAD) and mean relative deviation (MRD) for mass (M), center of mass (CoM), inertia (I), and kinetic energy (E). The best results are highlighted in bold. Inertia values only account for the diagonal elements of the inertia tensor, as generic OpenSim models assume off-diagonal elements to be zero.

| Model | Mean Absolute Deviation |  |  |  | Mean Relative Deviation |  |  |  |
| --- | --- | --- | --- | --- | --- | --- | --- | --- |
|  | M [g] | CoM [cm] | I [kg m<sup>2</sup>] | E [J] | M [%] | CoM [%] | I [%] | E [%] |
| scaled | 82.3 ± 28.4 | 8.7 ± 1.0 | 0.21 ± 0.42 | 2.1 ± 0.5 | 28.2 ± 8.6 | 60.4 ± 4.6 | 37.6 ± 15.1 | 26.4 ± 7.7 |
| SIPP | 56.6 ± 22.7 | **3.2 ± 2.2** | **0.17 ± 0.41** | 1.0 ± 0.9 | 10.2 ± 5.4 | **17.1 ± 7.8** | 16.3 ± 6.6 | 11.0 ± 7.7 |
| S-gen | 55.7 ± 53.9 | 3.4 ± 2.1 | 0.19 ± 0.40 | 1.0 ± 0.7 | **5.6 ± 4.9** | 18.4 ± 6.7 | 11.6 ± 5.0 | 9.9 ± 5.8 |
| addB | **33.1 ± 18.4** | 8.7 ± 1.0 | 0.22 ± 0.42 | 1.5 ± 0.4 | 22.1 ± 10.4 | 61.1 ± 5.1 | 32.6 ± 11.1 | 19.6 ± 5.8 |
| addB+SIPP | 53.9 ± 42.6 | 4.4 ± 2.8 | 0.19 ± 0.42 | 1.6 ± 0.8 | 17.1 ± 6.7 | 26.6 ± 19.6 | 23.3 ± 10.9 | 18.5 ± 4.9 |
| addB+S-gen | 91.8 ± 48.4 | 3.7 ± 2.5 | 0.18 ± 0.41 | **0.9 ± 0.4** | 5.7 ± 3.4 | 19.6 ± 9.0 | **3.7 ± 3.1** | **9.3 ± 2.7** |

While the effects of SIPP and AddBiomechanics’ optimization vary individually, a consistent trend emerges that all methods reallocate mass from distal to proximal lower limb segments, as illustrated in Fig. 2. The proximal mass reallocation is most pronounced for the SIPP model, and weakest for the AddBiomechanics physics optimization. In the SIPP and SIPP-generic models, the mass of the feet is approximately half of the mass in the scaled model.

**Fig 2.**
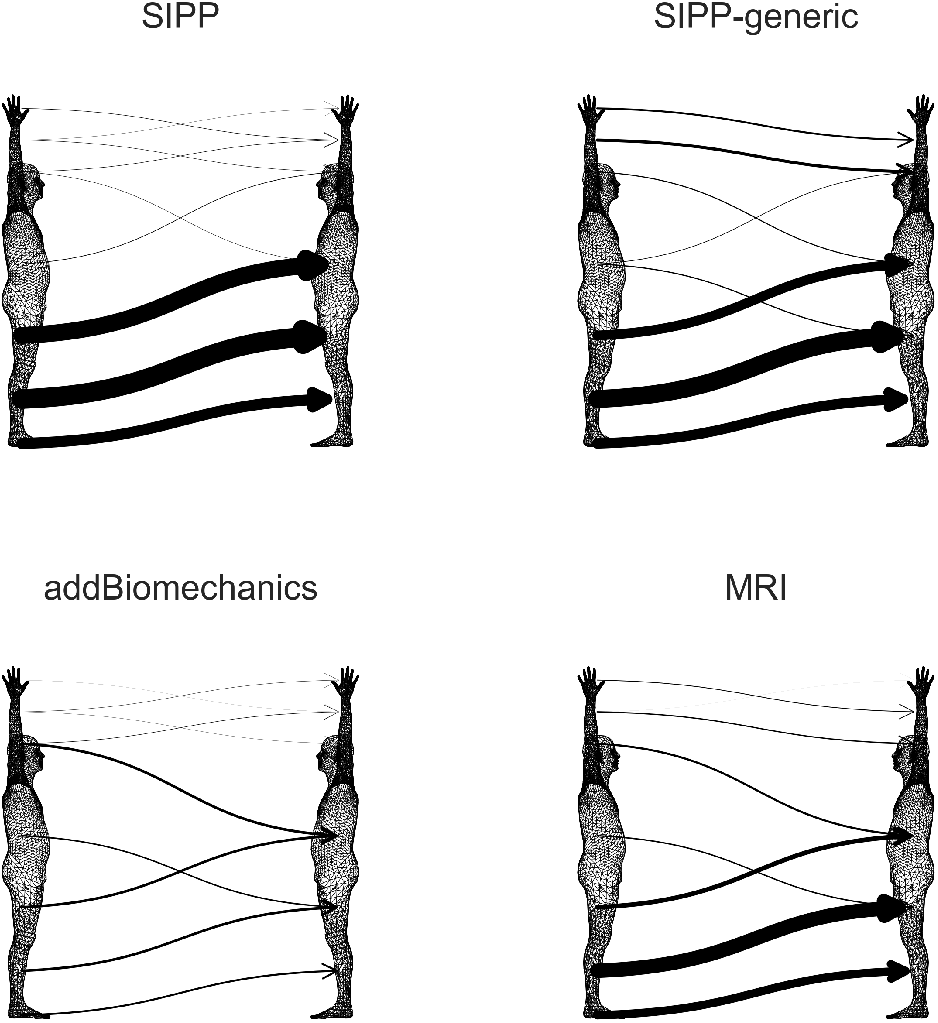
Average mass reallocation compared to scaled models. The arrows depict the average transfer of mass from the scaled (left) to the personalized MSK models (right). We binned the body parts into 7 regions, which are (from top to bottom): hand, lower arm, upper arm, trunk, upper leg, lower leg, and feet. The thickness of the arrows corresponds to the average amount of mass reallocated from one segment to another.

### Gait Analysis Outcomes

When using SIPP, residual forces reduced by 14.9 % and moments by 13.3 % compared to the scaled model. The MRI-personalized and SIPP-generic models did not affect the residuals noticeably, as shown in Fig. 3. After applying AddBiomechanics’ physics optimization, the residual forces and moments were reduced by 16.3 % and 13.2 % for SIPP, and 9.2 % and 4.2 % for SIPP-generic compared to a generic model optimized with AddBiomechanics’ physics optimization.

**Fig 3.**
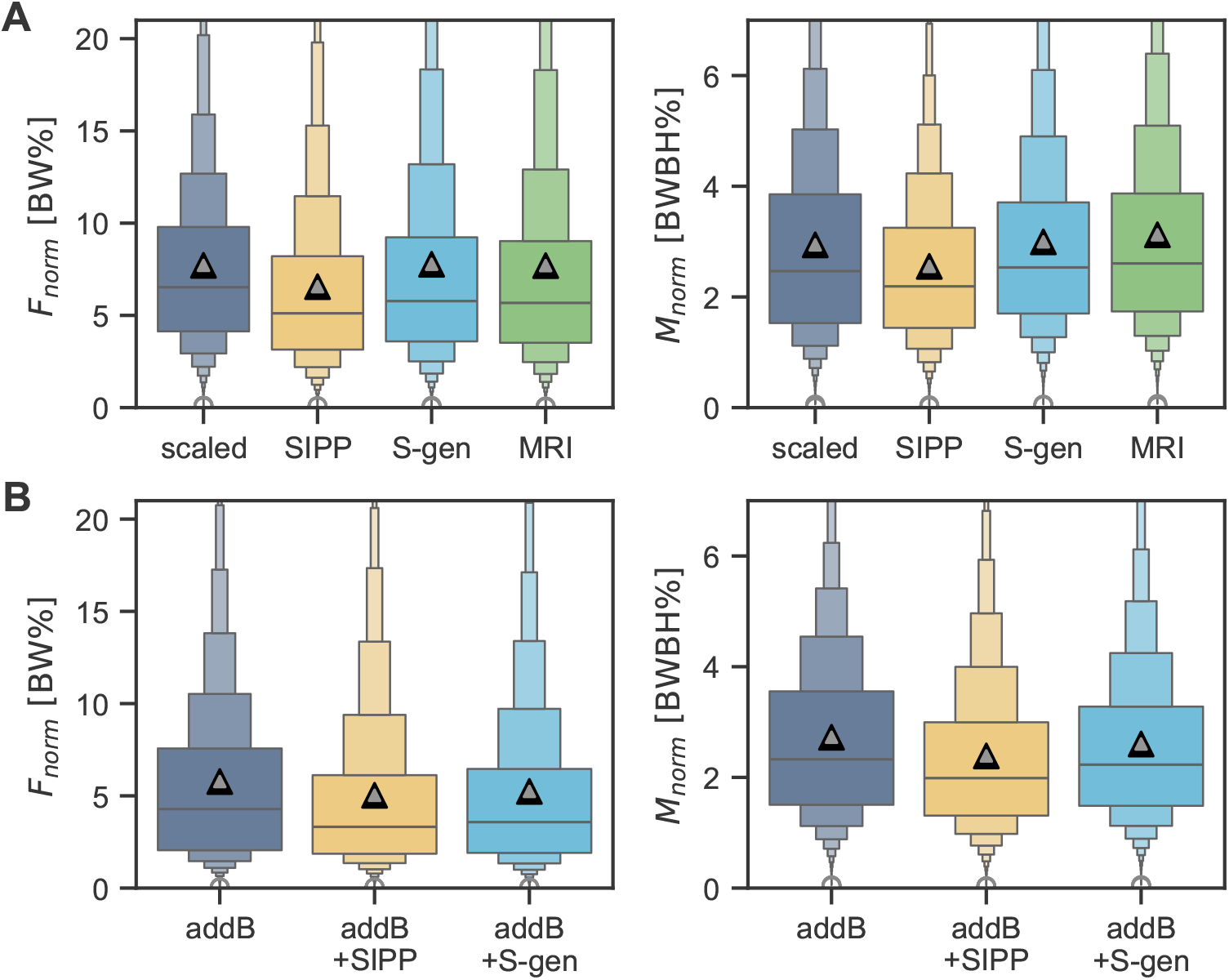
Residual forces and moments for all participants and trials. **A** shows the norm of the residual forces (left) and moments (right) for the MRI-personalized, scaled, SIPP, and SIPP-generic (S-gen) models. **B** shows the same residuals for the models in combination with AddBiomechanics’ physics optimization (addB, addB+SIPP, addB+S-gen). The residuals are shown as a percentage of bodyweight [BW%] for forces and as a percentage of bodyweight times body height [BWBH%] for moments. Means are indicated by grey triangles. Each box in the plot corresponds to progressively deeper quantile, e.g. the widest box covers the 50 % quantile, the next box covers the 75 % quantile, then the 87.5 % quantile, and so on. For visualization purposes, the top 5 % of data points (extreme outliers due to processing or measurement errors) are not shown in the y-axis range. Mean values were computed using the full dataset without exclusion.

Inverse dynamics joint moments were affected because the model’s BSIPs were altered, as shown in Fig. 4 and 5. In Fig. 4, we show the contralateral knee flexion moments for all participants, where a clear difference between the models can be seen during late swing phase, which is at 40 % to 50 % of the gait cycle. For 4 out of 6 participants, the mean values for the SIPP, SIPP-generic, and MRI-personalized models overlap during late swing phase, while the scaled model shows a larger moment. For the remaining two participants, the MRI-personalized model shows a larger moment than the SIPP and SIPP-generic models, and for one participant, the MRI-personalized model shows a moment similar to the scaled model. In Fig. 5, similar results can be seen for the hip and lumbar joints, where the SIPP, SIPP-generic, and MRI-personalized models all have lower joint moments than the scaled model. When using AddBiomechanics’ physics optimization (dashed lines), joint moment estimates change in a different way for each participant. For example, knee flexion moments during late swing phase are reduced for 5 out of 6 participants for the optimized generic model, but for participant 1, the optimized SIPP-generic model shows a larger moment than before optimization. On the other hand, the optimized SIPP model shows a lower moment than the unoptimized SIPP model for participant 1.

**Fig 4.**
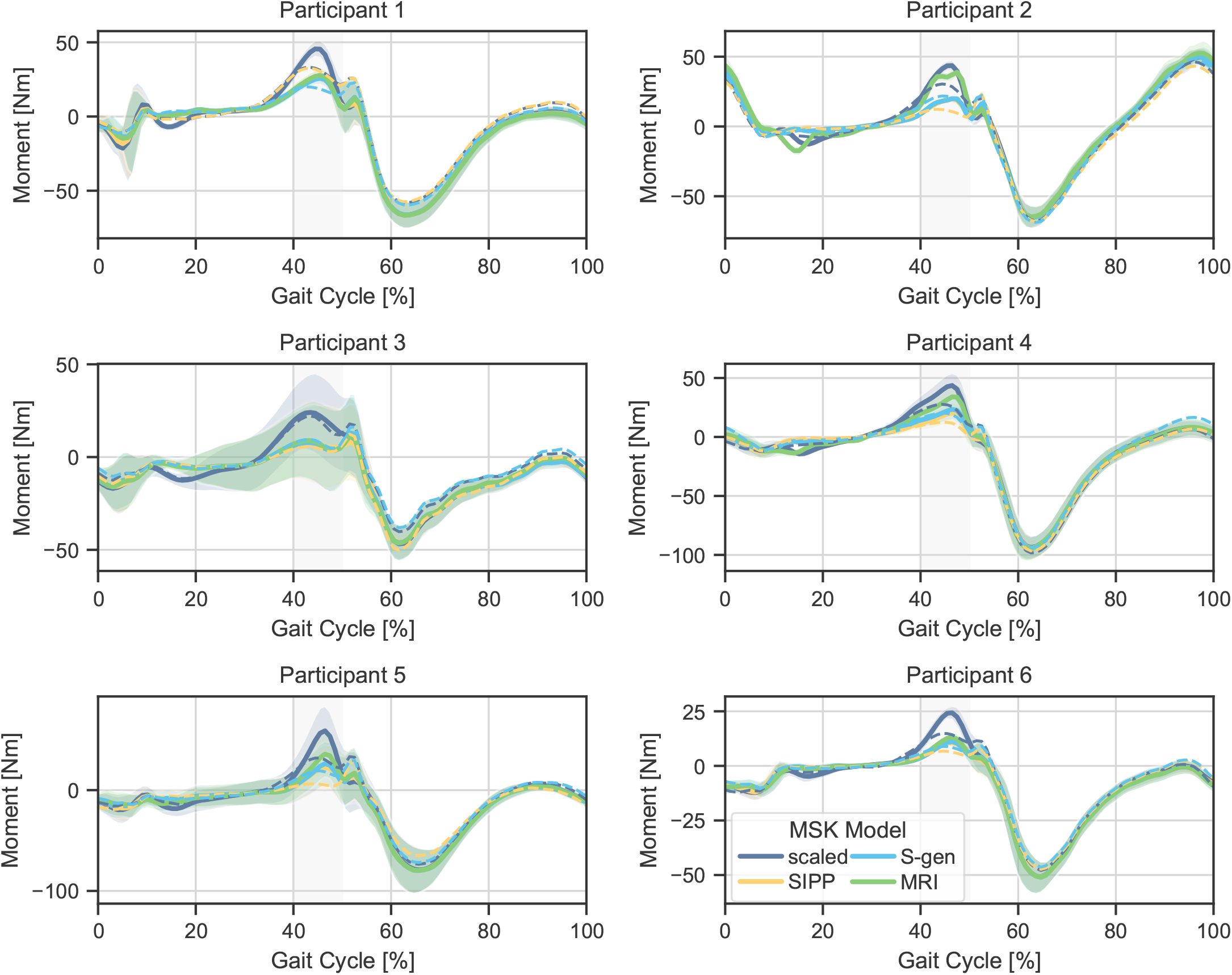
**Contralateral knee flexion moments for all participants during level walking at** 1.3 ms^*−*1^. Moments resulting from inverse dynamics are shown for the scaled, SIPP, SIPP-generic (S-gen), and MRI-personalized models. Shaded areas indicate standard deviation. Gait cycle lengths are normalized from heel strike to heel strike. We chose to show the contralateral knee, as the main difference between the models appears during late swing phase, which is at 40 to 50 % of the gait cycle (indicated by the gray-shaded regions). We also show models that are optimized with AddBiomechanics’ physics optimization as dashed lines.

**Fig 5.**
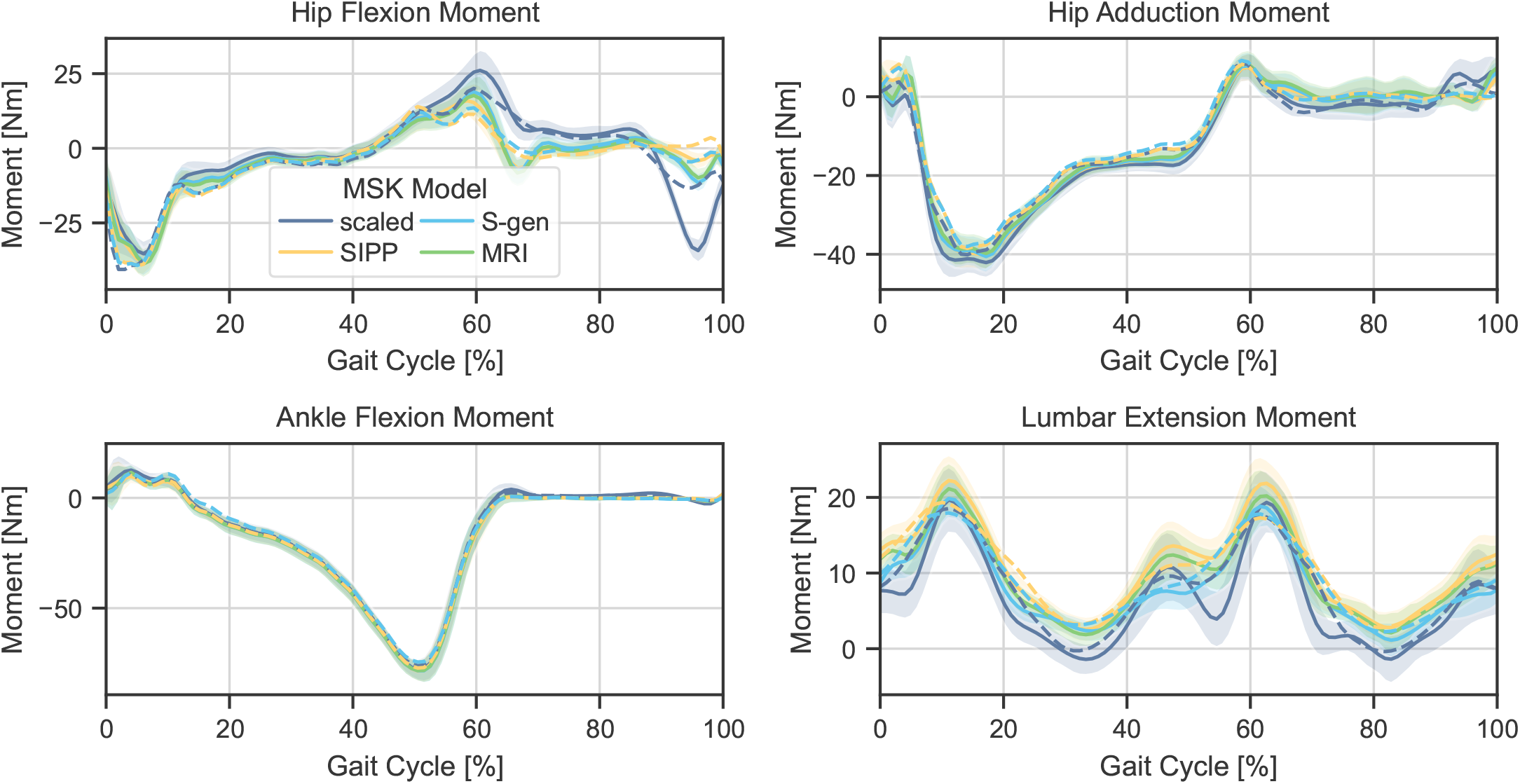
**Exemplary additional joint moments for one participant during level walking at** 1.3 ms^*−*1^. Moments resulting from inverse dynamics are shown for the scaled, SIPP, SIPP-generic (S-gen), and MRI-personalized models. Gait cycle lengths are normalized from heel strike to heel strike. Shaded areas indicate standard deviation. We also show models that are optimized with AddBiomechanics’ physics optimization as dashed lines.

By affecting joint moments, the choice of MSK model can lead to different metabolic cost estimates. Therefore, we calculated the metabolic costs according to Kim and Roberts’ joint space metabolic model [32]^1^. We observe that the mean metabolic cost estimation with SIPP is 7.5 % lower than that of the scaled model, and 12.8 % lower with SIPP+AddBiomechanics compared to AddBiomechanics alone. The MRI-personalized and SIPP-generic models show less difference in metabolic costs, with 1.7 % and 3.4 % lower costs than the scaled model, respectively. When using AddBiomechanics’ physics optimization, the metabolic costs for SIPP-generic and scaled models increase slightly, while the costs for SIPP-based models decrease. Metabolic cost estimates in different conditions are shown in Fig. 6.

**Fig 6.**
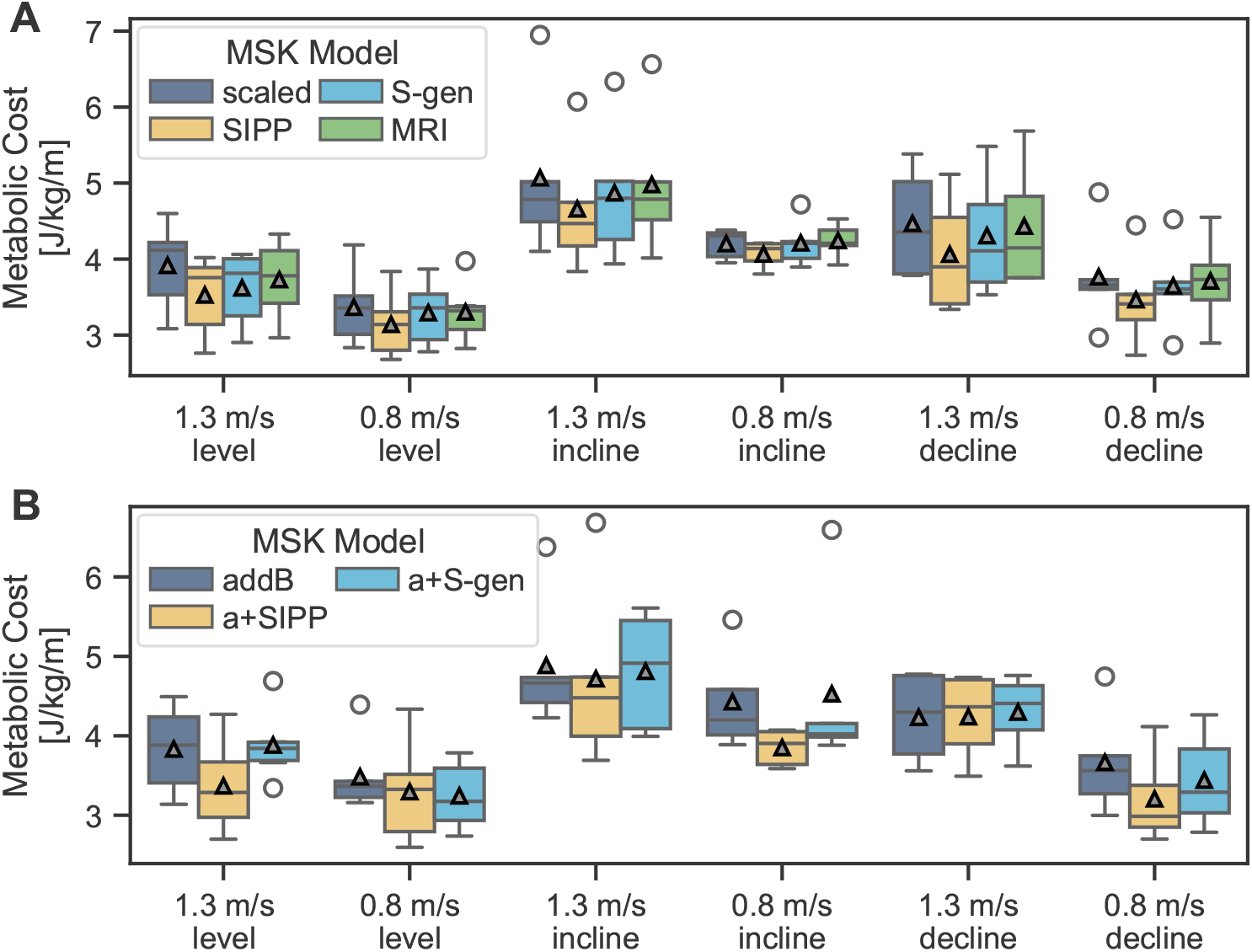
Metabolic costs per condition, as calculated with [32]. **A** shows the metabolic costs for the scaled, SIPP, SIPP-generic, and MRI-personalized models. **B** shows the same metabolic costs for the scaled, SIPP, and SIPP-generic models in combination with AddBiomechanics’ physics optimization (addB, a+SIPP, a+S-gen). Mean values are indicated by grey triangles. Outliers are indicated by unfilled circles.

### Sensitivity Study on Internal Tissue Estimation Model

In the sensitivity study on the internal tissue estimation model, we compared using SIPP using HIT, SIPP using InsideHumans, and SIPP without any internal tissue estimation. In the MRIgait benchmark, InsideHumans performed slightly better than HIT, but the differences were negligible. For gait-related results, however, SIPP with HIT outperformed both SIPP with InsideHumans and SIPP without internal tissue estimation. InsideHumans and no internal tissue estimation still led to reduced residual forces (InsideHumans: -5.4%, no internal tissue: -4.8%) and moments (InsideHumans: -4.3%, no internal tissue: -3.7%) compared to the scaled model, but the reductions were smaller than those achieved with HIT (-14.9% and -13.3%, respectively). Similarly, HIT reduced metabolic cost estimates (-7.5%) more than InsideHumans (-4.6%) and no internal tissue estimation (-5.4%). A detailed overview of the sensitivity study results is provided in S1 Appendix.

## Discussion

In this work, we introduced a method for shape-based inertial parameter personalization, SIPP, which we applied on easily acquirable smartphone pictures. We also applied SIPP on average body shapes to create, for the first time, gender-specific generic BSIP parameterizations. Furthermore, we introduced a benchmark dataset and task, on which we evaluated our method and compared it to a state of the art personalization method and MRI imaging. We showed that personalization of BSIPs impacts joint moments and that joint moments between MRI-based and SIPP-based BSIPs are similar, but differ from those estimated with current generic MSK models. As a consequence, metabolic costs are also reduced by up to 12.8 % when using SIPP. SIPP is also effective in reducing inverse dynamics residuals, which are a common problem in musculoskeletal modeling. Furthermore, we showed that SIPP-based BSIPs are in better agreement with the MRI-based BSIPs than the state of the art method, which is a major step towards personalized musculoskeletal modeling. Compared to MRI-based BSIP personalization, SIPP is more accessible as it requires no expert knowledge or time-intensive annotations.

In SIPP, we use HIT and SKEL [17, 18] to estimate the internal tissue distribution, as these are the highest-fidelity models available. To evaluate the sensitivity of SIPP towards the tissue estimation model, we tested two alternative pipelines, one where we skip the tissue estimation step, and another where we use InsideHumans instead of HIT [15]. While InsideHumans showed slightly more accurate BSIP than HIT on the MRIgait Benchmark, HIT showed a larger effect on residual forces and metabolics compared to when no tissue estimation model or InsideHumans was used. This highlights that the choice of internal tissue distribution model is crucial for accurate musculoskeletal modeling. Currently, HIT and SKEL are the best available models for biomechanical modeling. Nevertheless, these results also show that shape-based personalization of MSK models is generally effective, even when using lower-fidelity tissue distribution models. Inferring the internal tissue distribution from the body shape is ongoing research, and we expect that future models will provide more detailed and accurate tissue distributions, which will further improve the performance of SIPP.

Our findings are similar to those of Lee et al. [35], who used X-ray absorptiometry for BSIP estimation. They found that mass allocation to the foot and tibia was overestimated in current MSK models, and that late-swing phase joint moments were lower in their personalized model compared to various generic models. However, residual forces were not evaluated in their study. Due to the lack of practical applicability of X-ray absorptiometry, the method was never widely adopted. Personalization of BSIPs using less intrusive, image-based shape models has been proposed several times [36–38]; however, no method to date has been verified with medical imaging, shown to reduce inverse dynamics residuals, or reproduced Lee et al.’s findings on joint moments. Lee et al.’s finding on mass allocation has previously been confirmed by Chang et al. [36], although Chang’s study only included a single participant. Both Lee et al’.s and Chang et al.’s methods are not directly comparable to our method and dataset, as they either require different data modalities or do not provide an open-source pipeline for BSIP estimation.

In addition to direct estimation of a person’s BSIPs from their body, indirect methods exist that optimize the dynamic consistency of the model—such as AddBiomechanics or OpenSim’s residual reduction algorithm. To a lesser extent, AddBiomechanics tends to replicate trends observed in SIPP and by Lee et al., including mass shifting from the foot and thigh toward the femur and torso, and reduced swing-phase joint moments. However, these methods do not personalize the BSIPs but rather optimize the model to fit the data, which does not necessarily result in a more accurate representation of the individual’s body. While the reduction of joint moments through AddBiomechanics on generic models is consistent (see Fig. 4), its effect on personalized models can vary—either increasing or decreasing joint moments depending on the individual. Optimizing many BSIPs at once, with only six residual forces to fit, presents a non-linear optimization problem with many local minima, making initialization important. We found both SIPP and SIPP-generic to be suitable for initializing AddBiomechanics (see Table 2).

Current generic MSK models, such as Rajagopal2016, have more limited data origins and diversity than SIPP. In Rajagopal2016, Hamner’s model [39], and Arnold’s lower limb model [9], the BSIPs of the arms are based on gamma-ray scanning [12, 40], the torso and legs are based on photogrammetric measurements [11], and the feet are based on a low-fidelity 3D model of a size 10 tennis shoe from the 1990s [41]. These foundational studies were each conducted with less than 50 young, healthy, white male individuals, which is a very narrow demographic. In contrast, SIPP’s shape model, SMPL, is based on the CAESAR dataset [42], which includes 2.4k US/Canadian scans and 2k European scans, with half of the participants being female. The internal tissue distribution model, HIT, contains 241 female and 157 male MRI scans of unnamed origin [18]. While that gives SIPP and SIPP-generic a more diverse origin than current MSK models, it is still not representative of the general world-wide population, as non-western populations are not represented. We chose to not explicitly include a shoe model in SIPP, as the shoe mass varies between shoe types. This is a contributing factor to the mass reallocation as seen in Figure 2, where the foot mass is always higher in the generic model. We believe that the low-polygon model of a size 10 shoe, where the mass was set to a constant density of 1.1 g cm^*−*3^, is overestimating the foot mass, since convex features or pockets of air are neglected. Modern shoes are often made of lighter materials, which would further increase the discrepancy. Users of SIPP should be aware of this and add extra mass to the foot if they want to use SIPP for footwear research.

Humans naturally select gait patterns that minimize metabolic energy expenditure. The metabolic reduction observed from altering inertial parameters through SIPP of up to 12.8 % is relevant, as it is larger than the typical improvements seen with advanced footwear [43] and comparable to the gains achieved with modern exoskeleton assistance [44–46]. Therefore, we suggest that personalized modeling of BSIPs needs to be considered in exoskeleton and metabolics research. Another area where personalization can be relevant is in predictive simulations of human locomotion, where metabolic energy expenditure is often used as a cost function to optimize the simulation trajectory [47]. Often, knee range of motion during the swing phase is lower in simulations than in experiments [47, 48], which might be due to the higher joint moments that are needed to reproduce the experimental trajectory with a generic model. When using SIPP, the joint moments are lower, which might reflect a more optimal trajectory for the human body. Here, further studies are needed.

In processing of optical motion capture data, a lot of attention is given to low-pass filtering the data to reduce noise and artifacts [49]. The choice of the cutoff frequency impacts residual forces and joint moments [50]. Therefore, we also evaluated the sensitivity of SIPP’s improvement on residuals to the choice of cutoff frequency by running inverse dynamics on data filtered with cutoff frequencies between 6 Hz and 30 Hz. At 6 Hz, which is a common choice when analyzing gait data, residual forces and moments are lowest and the difference in residual forces is most pronounced, with SIPP causing a reduction of 24.2 % over the scaled model. The lowest effect on residual forces can be observed at 30 Hz, where SIPP reduces residual forces by 10.7 %. The absolute difference in residual forces between the scaled model and SIPP is relatively constant across all cutoff frequencies and varies between 1.1 BW% and 1.4 BW%. Residual moments are not greatly affected by the choice of cutoff frequency, with reductions between 13.1 % and 15.8 % for all cutoff frequencies. It has to be noted that the achievable lower bound for inverse-dynamics residuals fundamentally limited by measurement noise and rigid-body modeling assumptions, and that the improvements achieved with SIPP should be interpreted in this context.

The contributions of this paper - the MRIgait benchmark, smartphone-based body hull recording, and the SIPP-generic female musculoskeletal model - also pose ethical implications. Firstly, enabling body hull capture via smartphone is a significant step forward for MSK modeling in regions with limited healthcare infrastructure. This potential aligns with efforts to address equity in healthcare provision, whereby many countries and societies need to quickly achieve widespread healthcare provision. There is evidence that stakeholders expect low-cost technological solutions to achieve widespread coverage, reduce systemic healthcare disparities, and increase global healthcare justice [51]. In this context, some scholars argue for taking solidarity in account in the research, development, and distribution of new technologies [52, 53]. Secondly, the introduction of a female-generic SIPP model explicitly addresses unconscious bias in biomechanics [14] in the form of gender bias. It challenges the default use of male-derived models and refrains from framing female bodily variations as deviations. The SIPP-female-generic model marks an initial step towards more inclusive personalization. Firstly, because a gender-specific model makes it easier to identify and comprehend where (gender) bias occurs and influences simulations. As shown in the supporting document, scaled female MSK models perform worse on the MRIgait benchmark than their male counterparts. When using SIPP, on the other hand, female models perform better than male models on the benchmark, however, these findings should be interpreted cautiously, as the sample size is halved in gender-specific analyses.. A key difference between male and female SIPP and SIPP-generic models is the mass distribution between torso and thigh, where female MSK models have proportionally higher thigh mass than male MSK models. This difference in mass distribution is also reflected MRI-based MSK models, indicating that SIPP solves a relevant gender-specific aspect of BSIP personalization (see also S1 Appendix). Secondly, integrating the gender-balanced data sets provides initial indications of how discrimination along gender lines could be avoided in modeling. Future modeling and simulation can build on this by addressing other forms of bias, e.g. those related to aspects of bodily variation [54], and considering their intersection. An intersectional perspective is therefore essential to prevent discrimination based on physical deviations from prevailing body norms. In this respect, this study indicates how it might be possible to better represent deviations from the norm in future applications.

While SIPP is a step towards personalized musculoskeletal modeling, it has limitations, which are partially inherited from generic MSK models. For example, the geometry and muscle parameters of the Rajagopal2016 model are based on a 170 cm tall male person [10]. Furthermore, to take intersectional factors [55, 56] into account, and further diversification across age, body types, or different gender identities, could improve representation and applicability. We avoided tight-fitting body hulls due to practical challenges such as clothing artifacts and processing complexity, while we found that the parametric hull typically provides a sufficient fit. Recently, length-based scaling based on body hulls has been explored [20], and muscle parameters may correlate with body hull parameterizations [57], suggesting potential for complete musculoskeletal personalization using body hulls when combined with SIPP. Additionally, while HIT provides the best performance, a more granular tissue estimation model that can, e.g., seperate abdominal muscles from organs, or cortical from trabecular bone, could further enhance BSIP accuracy, and we expect SIPP to improve with future internal tissue distribution models, as well as integration with retrospective personalization methods [6, 58]. We used smartphone pictures for SIPP because they are quick to acquire and widely available, to allow any researcher to benefit from SIPP. To that end, we open-source our implementation and data (see S3 Code Repository and S4 Data Repository). Currently, technical expertise is required to set up multiple Python environments for data processing. We aim to improve on this in the future by releasing a web-app that automates the data processing completely.

However, for clinical applications, clinically approved 3D body scanners should be used, as they provide a more standardized procedure for body hull acquisition and reliable reconstructions of the upper limbs.

## Conclusion

We presented a method to personalize inertial parameters of musculoskeletal models based on smartphone-recorded body hulls. We validated our method on a small cohort of participants, where we also acquired MRI scans and gait data. Our results show that our method leads to reduced residual forces and joint torques, as well as lower metabolic cost estimations when compared to generic musculoskeletal models. Furthermore, we introduced two new generic musculoskeletal models that are based on average body shapes and can be used when no body hulls are available. We also introduced the MRIgait benchmark, which allows to compare inertial parameter personalization methods based on their deviations from MRI-generated MSK models. Our research makes personalized musculoskeletal models more accessible and supports broader use in research and, when using validated body scanners, clinical settings.

## Supporting information

### S1 Appendix. Supporting document

This document contains additional information on the participant demographics and additional results.

### S2 Project Page

A project website linking to all relevant repositories and data is available at https://gambimar.github.io/sipp/.

### S3 Code Repository

The code for the SIPP method, the MRIgait benchmark and the SIPP-generic models is available at https://github.com/gambimar/sipp.

### S4 Data Repository

The MRIgait dataset, results, and MRIgait benchmark repository are available at https://doi.org/10.5281/zenodo.15791380.

### S5 SimTK Project

We made SIPP-generic available to the OpenSim community via SimTK at https://simtk.org/projects/sipp-generic.

## Acknowledgments

This work was funded by the Deutsche Forschungsgemeinschaft (DFG, German Research Foundation) – SFB 1483 – Project-ID 442419336, EmpkinS. This work was also supported with an International Travel Grant by the International Society of Biomechanics (ISB).

## Author Contributions

**Conceptualization:** Markus Gambietz, Iris Wechsler, Jörg Miehling, Mario Botsch, Anne D. Koelewijn **Methodology:** Markus Gambietz, Iris Wechsler, Philipp Amon, Mario Botsch, Timo Menzel, Katie McMahon, Anne D. Koelewijn **Software:** Markus Gambietz, Philipp Amon, Timo Menzel, Mario Botsch **Validation:** Markus Gambietz **Formal analysis:** Markus Gambietz **Investigation:** Markus Gambietz **Data Curation:** Markus Gambietz, Philipp Amon, Putri Qistina Azam **Writing - Original Draft:** Markus Gambietz, Eva Maria Hille, Tabea Ott, Matthias Braun **Writing - Review & Editing:** Markus Gambietz, Iris Wechsler, Eva Maria Hille, Tabea Ott, Matthias Braun, Jörg Miehling, Mario Botsch, Katie McMahon, Anne D. Koelewijn **Visualization:** Markus Gambietz **Supervision:** Katie McMahon, Jörg Miehling, Anne D. Koelewijn **Project administration:** Anne D. Koelewijn **Funding acquisition:** Katie McMahon, Anne D. Koelewijn

## Footnotes

1 The choice of metabolic model is based on its applicability on inverse dynamics outputs and its high correlation with experimental data [33, 34].

## Notes

https://github.com/gambimar/smartphone_personalization_mri

https://simtk.org/projects/sipp-generic

https://zenodo.org/records/15791380

### Competing Interest Statement

The authors have declared no competing interest.

### Summary of Updates

Title and abstract revised to clarify contributions and reduce emphasis on smartphone imaging. Interpolation method corrected (thin-plate spline → cubic kernel); results updated without affecting conclusions. Introduction and Methods clarified, including definitions of key models, improved reproducibility of the SIPP pipeline, runtime (~5 min), and data acquisition details. Results and figures corrected (cross-references, captions, outlier handling) with additional analyses moved to supplementary material. Discussion expanded to address reviewer concerns (e.g., sex-specific trends, model comparisons, limitations). Language, formatting, and hyperlinks standardized; supplementary files updated with additional quantitative results.

