## Supplementary material for "From Body Hulls to Musculoskeletal Models: Personalized Inertial Parameter Estimation": S1 Appendix

### Supporting Information for: From Smartphone Images to Musculoskeletal Models: Personalized Inertial Parameter Estimation

In this supporting document, we provide additional results and details that support the findings presented in the main text. This includes participant and trial metadata, additional results for joint moments, and a detailed analysis the influence of gender on personalization and gait analysis outcomes.

#### Participant and Trial Metadata

Participant and trial information are summarized in Table A.1 and Table A.2, respectively. Body height, age and gender are self-reported, while body mass was measured using force plates. The excluded trials are: P01, trials 3-6 and P06, trial 5.

Table A.1: **Participant Information.**

| ID | Age (years) | Mass (kg) | Height (cm) | Sex |
|---|---|---|---|---|
| P01 | 26 | 72.31 | 167 | F |
| P02 | 28 | 73.35 | 177 | M |
| P03 | 26 | 64.49 | 165 | F |
| P04 | 27 | 89.36 | 177 | M |
| P05 | 30 | 87.63 | 179 | M |
| P06 | 26 | 44.91 | 147 | F |

Table A.2: **Trial Information.**

| Trial Number | Speed (m/s) | Incline (%) |
|---|---|---|
| 1 | 1.3 | 0 |
| 2 | 0.8 | 0 |
| 3 | 1.3 | 8 |
| 4 | 0.8 | 8 |
| 5 | 1.3 | -8 |
| 6 | 0.8 | -8 |

#### Additional Results

Similar to Fig. 4 in the main text, we present the joint moments for all models for lumbar rotation A.1, lumbar bending A.2, lumbar extension A.3, hip flexion A.4, hip rotation A.5, hip adduction A.6, and ankle angle A.7. Next, in Table A.3, we present the residual forces for each model.

The sensitivity of SIPP towards different internal tissue models are included in Tables A.3 and A.4, and metabolics for these pipelines are shown in Fig. A.8. The nominal SIPP variant uses Human Implicit Tissues (HIT) [1] to infer internal tissue distributions. We also show results for SIPP using Komaritzan et al.'s layered tissue model "InsideHumans" (IH) [2] and SIPP without internal tissue estimations (w/o HIT). Tissue estimation methods have only minor influence on the MRIgait Benchmark results, with SIPP (IH) showing slightly better performance than the others. On the other hand, nominal SIPP using HIT shows slightly better residual forces (SIPP: -14.9%; SIPP (IH): -4.8%; SIPP (w/o HIT): -5.4%) and slightly better residual moments (SIPP: -13.3%; SIPP (IH): -11.2%; SIPP (w/o HIT): -8.8% relative to the scaled model). Metabolic costs are also lower for nominal SIPP compared to the other two versions (SIPP: -7.6%; SIPP (IH): -4.6%; SIPP (w/o HIT): -5.4% relative to the scaled model).

Table A.3: **Overview of residual forces.** We show the mean and standard deviation of the absolute residuals for all dimensions ($x$, $y$, $z$) and their norms. Forces are listed as a percentage of bodyweight [BW%] and the values for absolute residual moments are listed as a percentage of bodyweight times bodyweight [BWBH%]. We show the results for the MRI-personalized model, scaled model, SIPP model, and SIPP-generic (S-gen) model in the first six rows. In addition to the nominal SIPP variant using Human Implicit Tissues (HIT) [1] to infer internal tissue distributions, we also show results for SIPP using Komaritzan et al.'s layered tissue model "InsideHumans" (IH) [2] and SIPP without internal tissue estimations (w/o HIT). In the last three rows, we show the results for the same models in combination with addBiomechanics' physics optimization (addB, add+SIPP, add+S-gen). The best results per table section are highlighted in bold.

| | **Forces** | | | | **Moments** | | | |
|---|---|---|---|---|---|---|---|---|
| **Model** | $\mathbf{F}_x$ [BW%] | $\mathbf{F}_y$ [BW%] | $\mathbf{F}_z$ [BW%] | $\|\mathbf{F}\|$ [BW%] | $\mathbf{M}_x$ [BWBH%] | $\mathbf{M}_y$ [BWBH%] | $\mathbf{M}_z$ [BWBH%] | $\|\mathbf{M}\|$ [BWBH%] |
| MRI | $4.75 \pm 7.0$ | $3.50 \pm 3.9$ | $3.04 \pm 3.2$ | $7.70 \pm 7.7$ | $1.75 \pm 1.8$ | $0.92 \pm 0.8$ | $1.91 \pm 4.8$ | $3.12 \pm 5.0$ |
| scaled | $5.14 \pm 5.1$ | $3.57 \pm 3.2$ | $\mathbf{2.67 \pm 2.7}$ | $7.71 \pm 5.5$ | $2.06 \pm 1.8$ | $0.96 \pm 0.8$ | $\mathbf{1.30 \pm 4.3}$ | $2.94 \pm 4.5$ |
| SIPP | $\mathbf{3.90 \pm 4.8}$ | $\mathbf{3.07 \pm 3.2}$ | $2.73 \pm 2.7$ | $\mathbf{6.56 \pm 5.5}$ | $\mathbf{1.43 \pm 1.3}$ | $\mathbf{0.86 \pm 0.7}$ | $1.49 \pm 4.3$ | $\mathbf{2.55 \pm 4.4}$ |
| SIPP (IH) | $4.45 \pm 6.5$ | $3.40 \pm 3.9$ | $2.94 \pm 3.1$ | $7.34 \pm 7.2$ | $1.59 \pm 1.7$ | $0.87 \pm 0.8$ | $1.39 \pm 4.7$ | $2.61 \pm 4.9$ |
| SIPP (w/o HIT) | $4.40 \pm 6.2$ | $3.38 \pm 3.9$ | $2.94 \pm 3.1$ | $7.29 \pm 7.0$ | $1.57 \pm 1.6$ | $0.88 \pm 0.8$ | $1.52 \pm 4.8$ | $2.68 \pm 4.9$ |
| S-gen | $4.66 \pm 6.8$ | $3.56 \pm 4.0$ | $3.19 \pm 3.4$ | $7.77 \pm 7.6$ | $1.64 \pm 1.7$ | $0.92 \pm 0.8$ | $1.83 \pm 4.8$ | $3.00 \pm 4.9$ |
| addB | $3.51 \pm 5.7$ | $3.38 \pm 4.6$ | $\mathbf{1.63 \pm 2.9}$ | $5.77 \pm 7.4$ | $1.85 \pm 1.5$ | $0.93 \pm 0.8$ | $\mathbf{1.25 \pm 4.7}$ | $2.72 \pm 4.9$ |
| addB+SIPP | $\mathbf{2.62 \pm 4.0}$ | $\mathbf{2.84 \pm 3.7}$ | $1.69 \pm 2.8$ | $\mathbf{4.83 \pm 5.6}$ | $\mathbf{1.29 \pm 1.2}$ | $\mathbf{0.86 \pm 0.7}$ | $1.35 \pm 4.3$ | $\mathbf{2.34 \pm 4.3}$ |
| addB+S-gen | $2.79 \pm 4.9$ | $3.18 \pm 4.6$ | $1.77 \pm 3.9$ | $5.24 \pm 7.3$ | $1.40 \pm 1.2$ | $0.89 \pm 0.8$ | $1.60 \pm 4.8$ | $2.60 \pm 4.8$ |

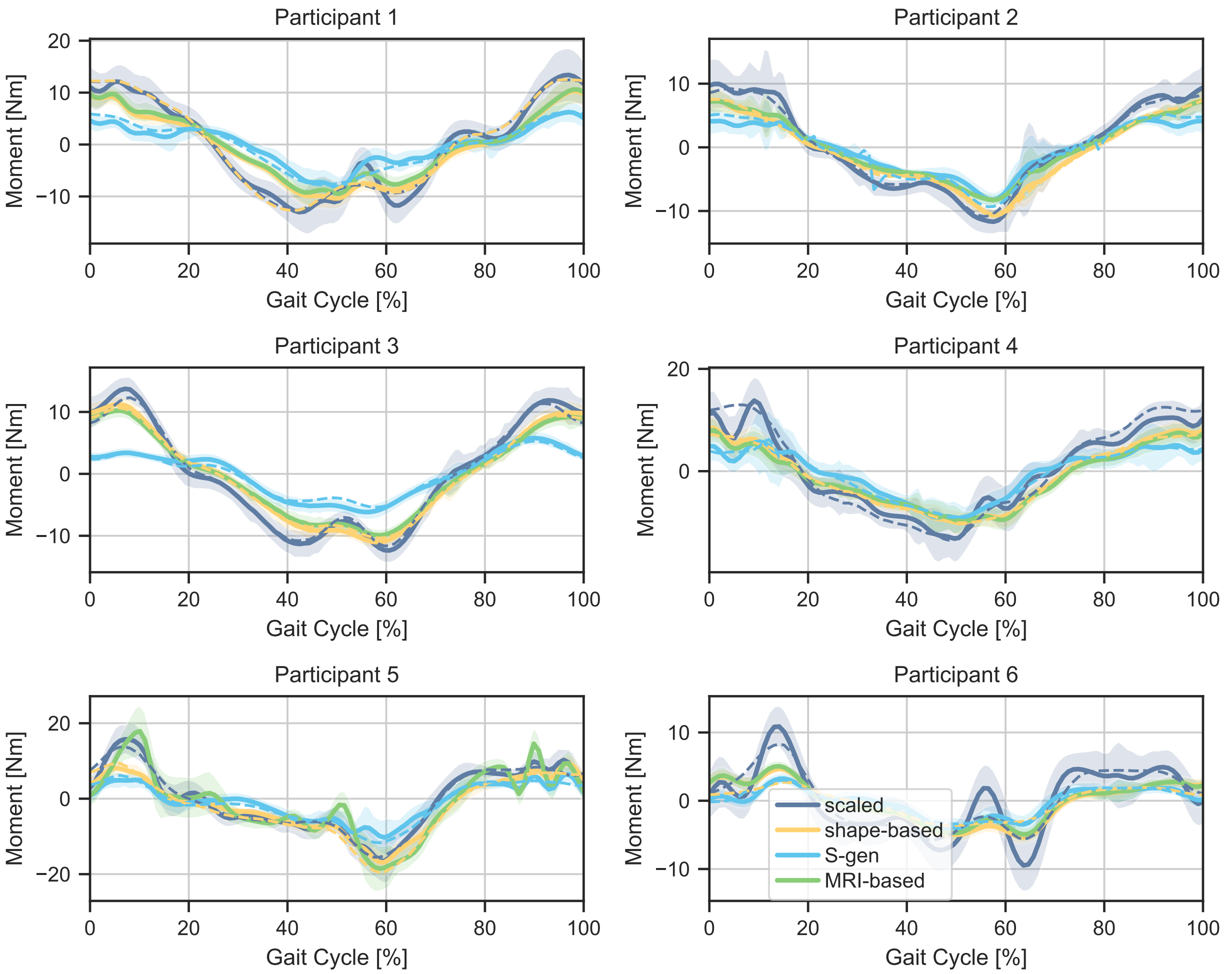


Figure A.1: **Lumbar rotation moments for all participants during level walking at $1.3\,\mathrm{m\,s^{-1}}$.** Moments resulting from inverse dynamics are shown for the scaled, SIPP, SIPP-generic (S-gen), and MRI-personalized models. Shaded areas indicate standard deviation. We also show models that are optimized with addBiomechanics' physics optimization as dashed lines.

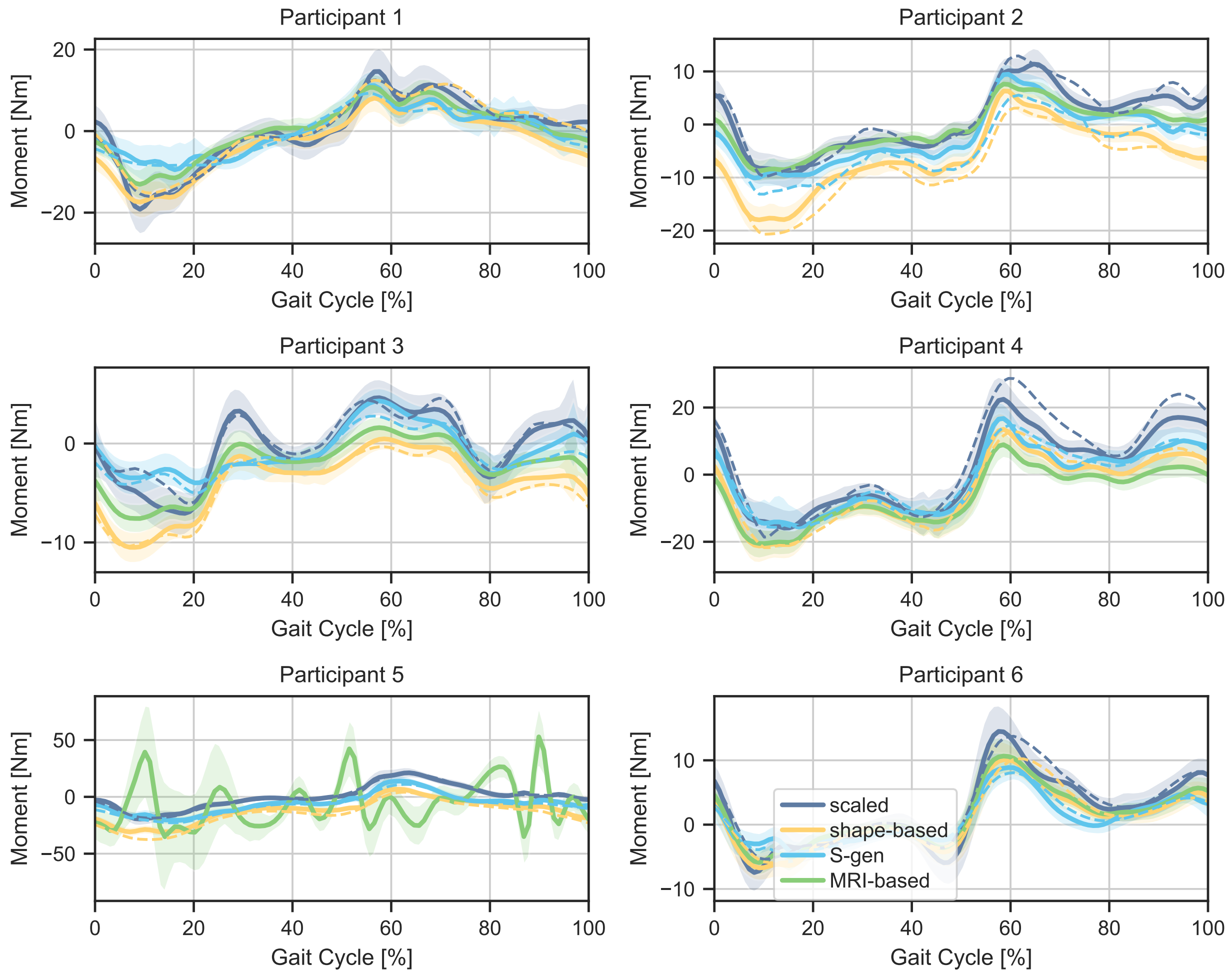


Figure A.2: **Lumbar bending moments for all participants during level walking at $1.3\,\mathrm{m\,s^{-1}}$.** Moments resulting from inverse dynamics are shown for the scaled, SIPP, SIPP-generic (S-gen), and MRI-personalized models. Shaded areas indicate standard deviation. We also show models that are optimized with addBiomechanics' physics optimization as dashed lines.

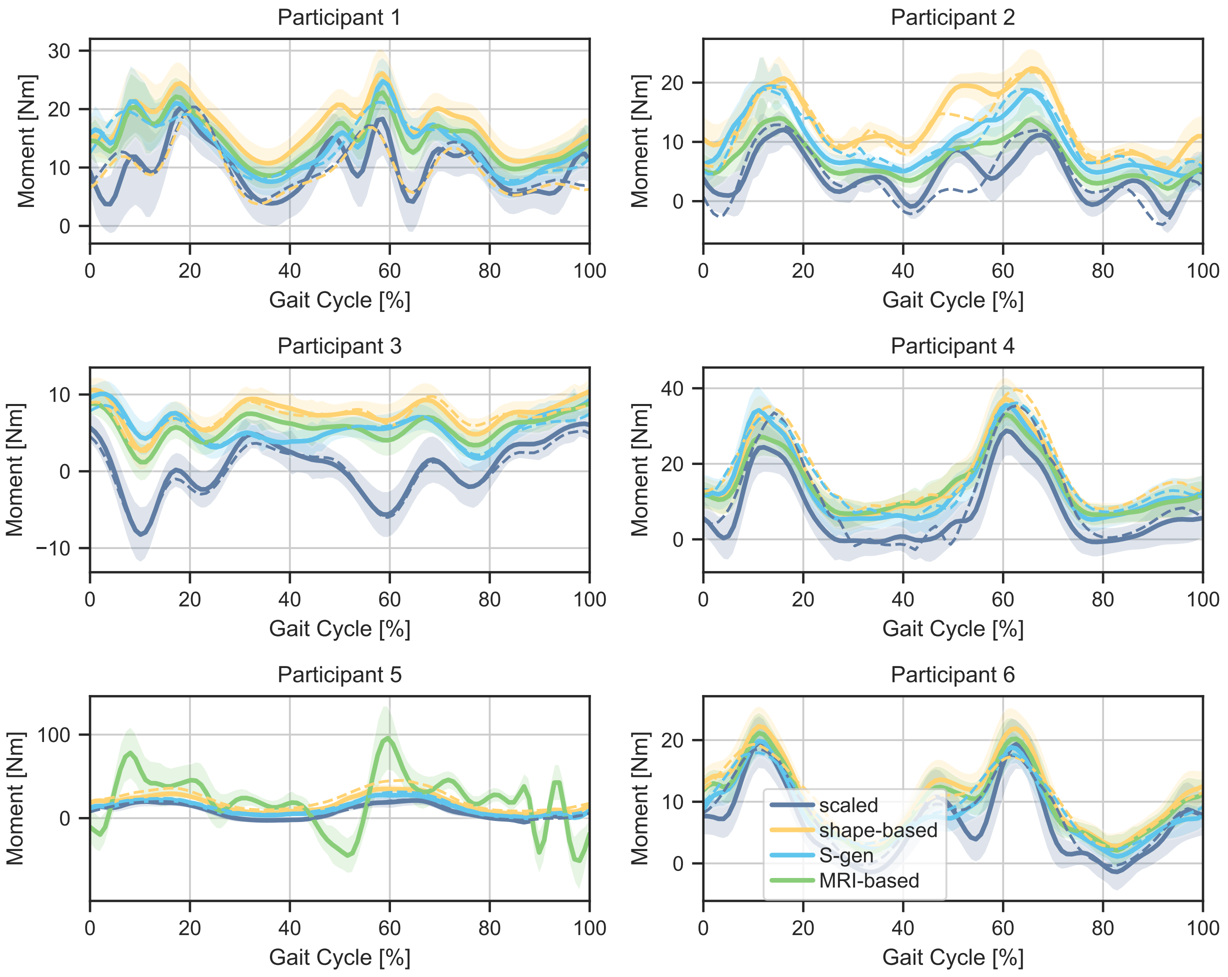


Figure A.3: **Lumbar extension moments for all participants during level walking at $1.3\,\mathrm{m\,s^{-1}}$.** Moments resulting from inverse dynamics are shown for the scaled, SIPP, SIPP-generic (S-gen), and MRI-personalized models. Shaded areas indicate standard deviation. We also show models that are optimized with addBiomechanics' physics optimization as dashed lines.

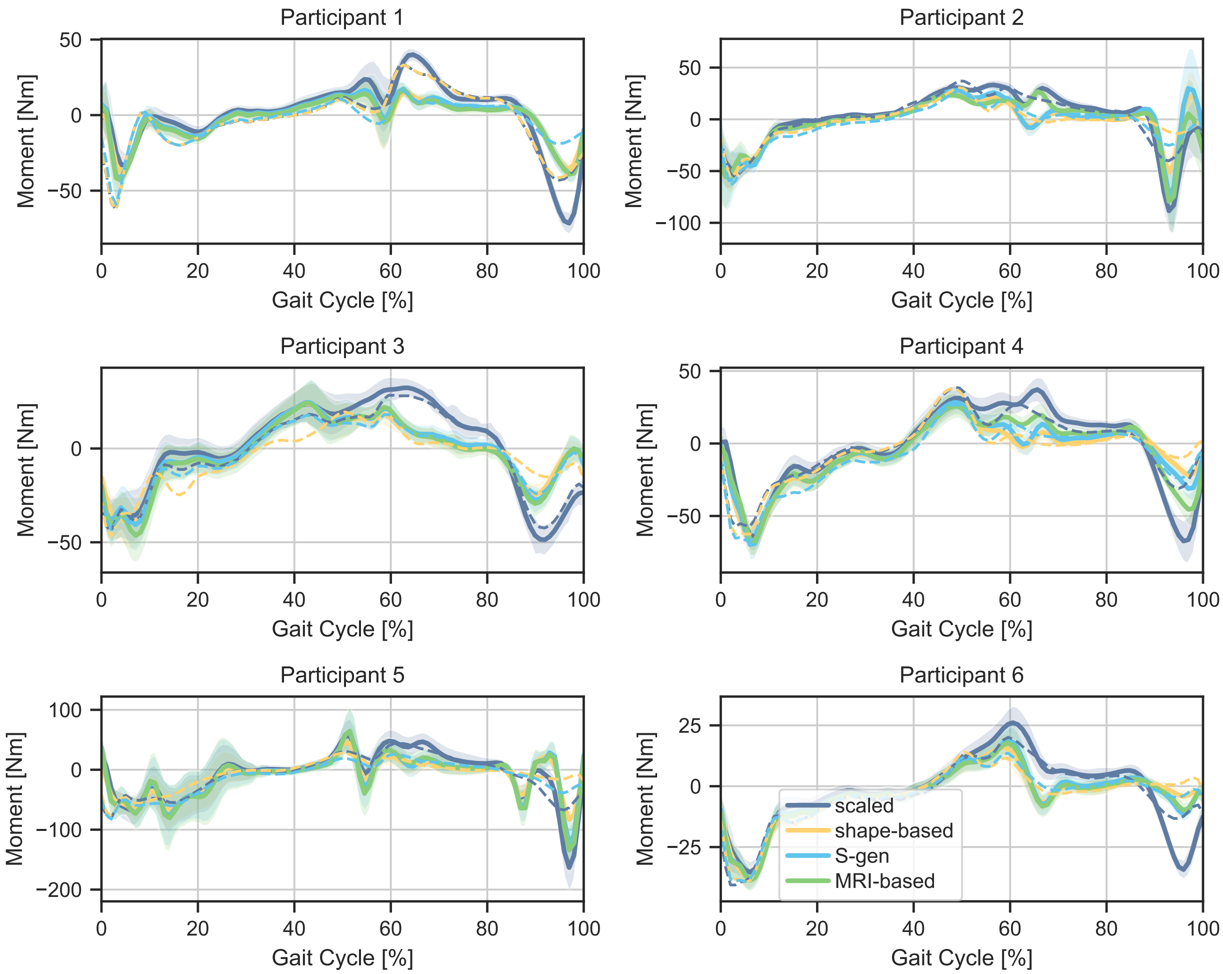


Figure A.4: **Hip flexion moments for all participants during level walking at $1.3\,\mathrm{m\,s^{-1}}$.** Moments resulting from inverse dynamics are shown for the scaled, SIPP, SIPP-generic (S-gen), and MRI-personalized models. Shaded areas indicate standard deviation. We also show models that are optimized with addBiomechanics' physics optimization as dashed lines.

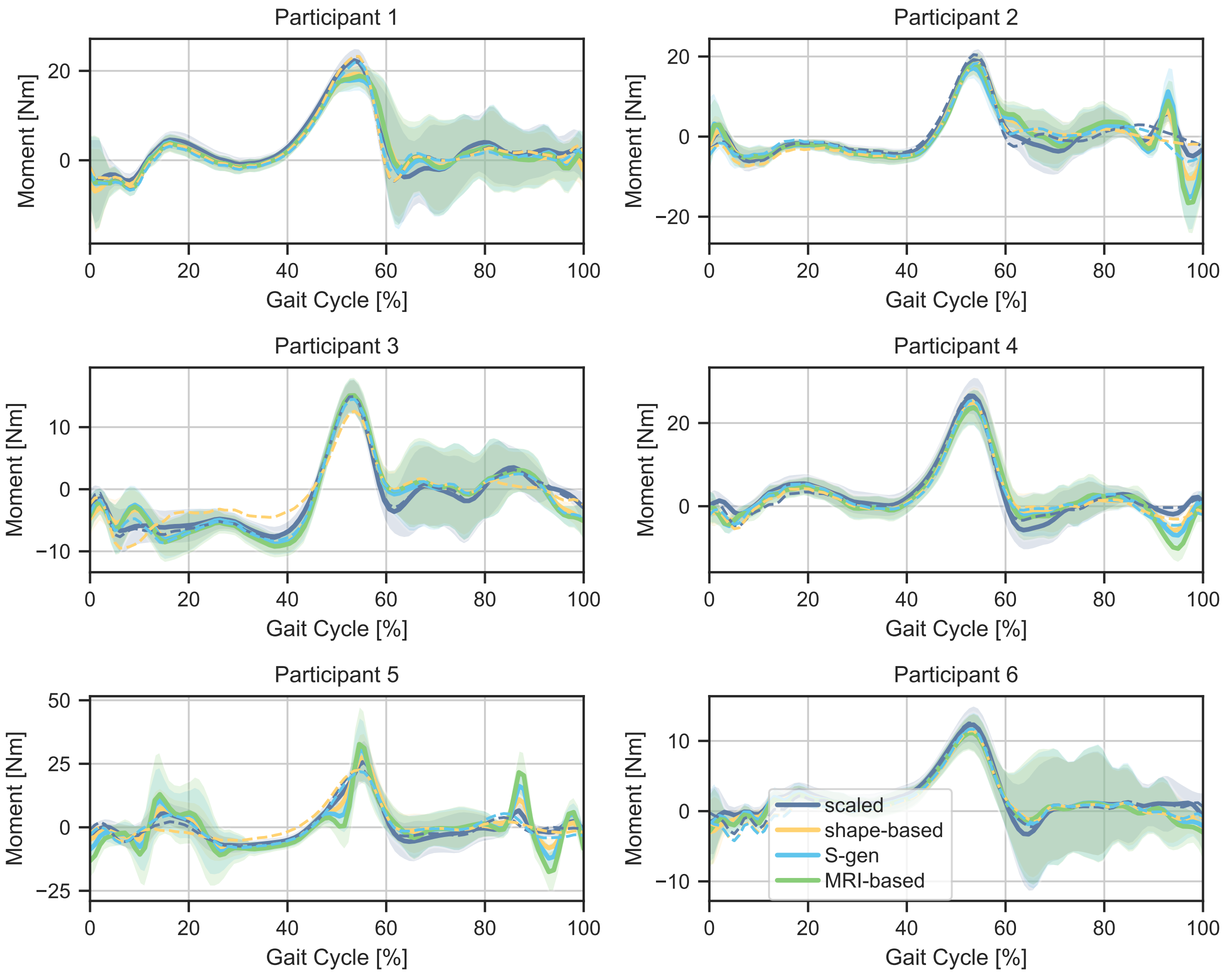


Figure A.5: **Hip rotation moments for all participants during level walking at $1.3\,\mathrm{m\,s^{-1}}$.** Moments resulting from inverse dynamics are shown for the scaled, SIPP, SIPP-generic (S-gen), and MRI-personalized models. Shaded areas indicate standard deviation. We also show models that are optimized with addBiomechanics' physics optimization as dashed lines.

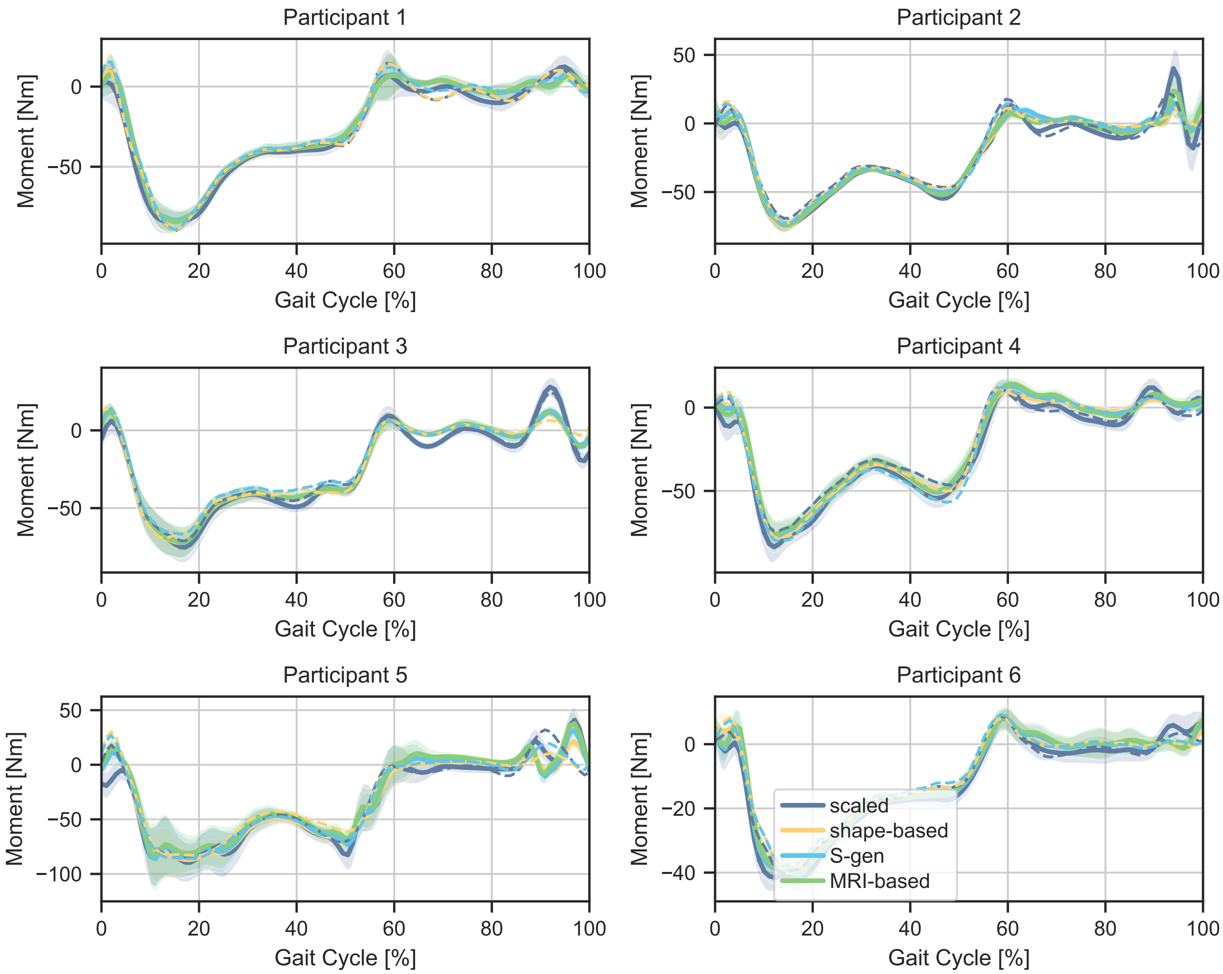


Figure A.6: **Hip adduction moments for all participants during level walking at $1.3\,\mathrm{m\,s^{-1}}$.** Moments resulting from inverse dynamics are shown for the scaled, SIPP, SIPP-generic (S-gen), and MRI-personalized models. Shaded areas indicate standard deviation. We also show models that are optimized with addBiomechanics' physics optimization as dashed lines.

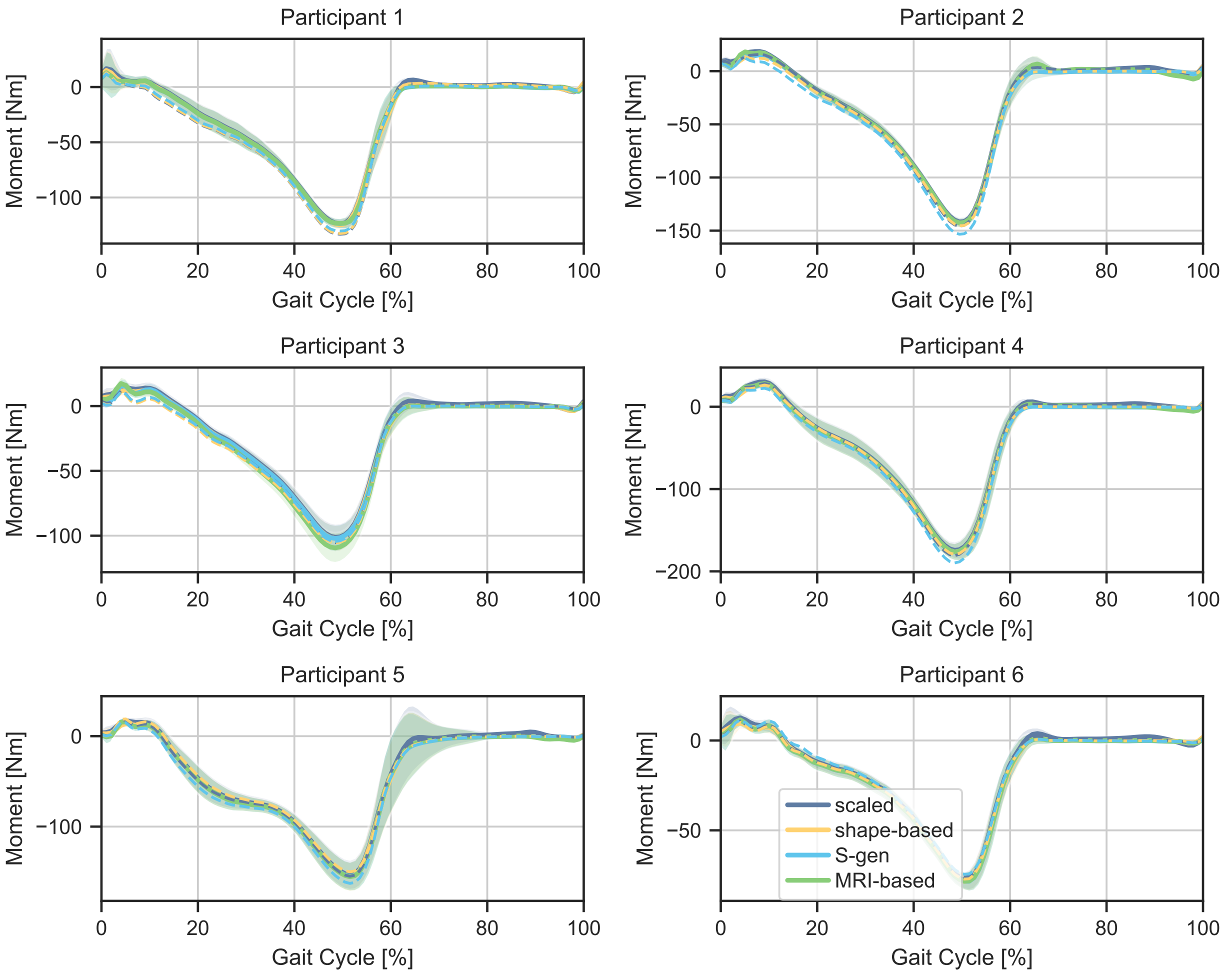


Figure A.7: **Ankle moments for all participants during level walking at** $1.3\,\mathrm{m\,s^{-1}}$**.** Moments resulting from inverse dynamics are shown for the scaled, SIPP, SIPP-generic (S-gen), and MRI-personalized models. Shaded areas indicate standard deviation. We also show models that are optimized with addBiomechanics' physics optimization as dashed lines.

Table A.4: **Internal tissue estimation models on the MRIgait Benchmark.** This table shows results for SIPP using different internal tissue models. The first row shows the nominal SIPP variant using Human Implicit Tissues (HIT) [1] to infer internal tissue distributions. The second row shows results for SIPP using Komaritzan et al.'s layered tissue model "InsideHumans" (IH) [2], and the third row shows results for SIPP without internal tissue estimations (w/o HIT). The columns show the mean and standard deviation of the mean absolute deviation (MAD) and mean relative deviation (MRD) for mass (M), center of mass (CoM), inertia (I), and kinetic energy (E). The best results are highlighted in bold. Inertia values only account for the diagonal elements of the inertia tensor, as generic OpenSim models assume off-diagonal elements to be zero.

| | **Mean Absolute Deviation** | | | | **Mean Relative Deviation** | | | |
|---|---|---|---|---|---|---|---|---|
| **Model** | **M** [g] | **CoM** [cm] | **I** [kg m$^2$] | **E** [J] | **M** [%] | **CoM** [%] | **I** [%] | **E** [%] |
| scaled | $82.3 \pm 28.4$ | $8.7 \pm 1.0$ | $0.21 \pm 0.42$ | $2.1 \pm 0.5$ | $28.2 \pm 8.6$ | $60.4 \pm 4.6$ | $37.6 \pm 15.1$ | $26.4 \pm 7.7$ |
| SIPP | $56.6 \pm 22.7$ | $\mathbf{3.2 \pm 2.2}$ | $0.17 \pm 0.41$ | $1.0 \pm 0.9$ | $10.2 \pm 5.4$ | $17.1 \pm 7.8$ | $16.3 \pm 6.6$ | $11.0 \pm 7.7$ |
| SIPP (IH) | $\mathbf{55.7 \pm 22.0}$ | $3.2 \pm 2.2$ | $\mathbf{0.17 \pm 0.41}$ | $\mathbf{0.9 \pm 0.8}$ | $\mathbf{9.6 \pm 4.6}$ | $\mathbf{17.0 \pm 7.6}$ | $\mathbf{12.1 \pm 6.2}$ | $\mathbf{9.8 \pm 7.1}$ |
| SIPP (w/o HIT) | $57.1 \pm 22.6$ | $3.2 \pm 2.2$ | $0.17 \pm 0.41$ | $1.0 \pm 0.9$ | $9.9 \pm 5.1$ | $17.2 \pm 7.8$ | $14.7 \pm 6.9$ | $10.7 \pm 7.6$ |

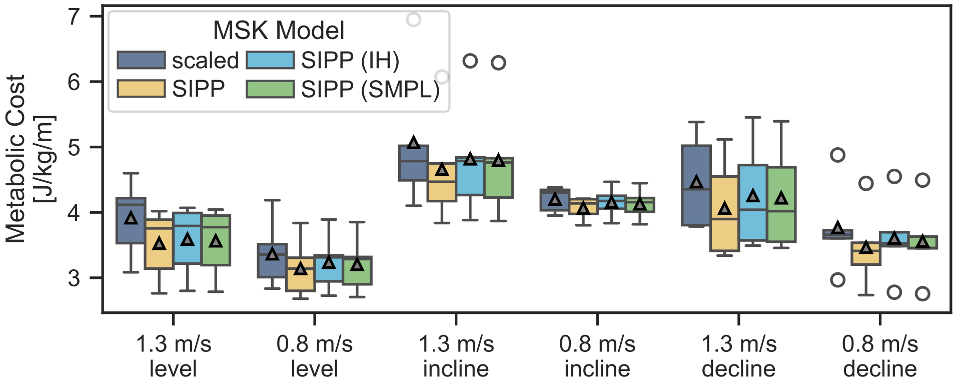


Figure A.8: **Metabolic costs per condition, as calculated with [3].** We show the scaled model compared to SIPP and alternative versions using either Komaritzan et al.'s layered tissue model "InsideHumans" (IH) [2] or no internal tissue estimations (w/o HIT). Mean values are indicated by grey triangles. Outliers are indicated by unfilled circles.

### Effect of gender on SIPP-generic

#### Results per gender

Female MSK models perform worse on all mean relative deviations in the MRIgait benchmark than scaled male MSK models, as shown in Table A.5. Both SIPP and SIPP-generic models outperform the scaled models for each gender. Mean absolute deviations should not be compared between genders, as the body mass and height greatly differ. In Figs A.9, we show the average mass reallocation when using SIPP, SIPP-generic, addBiomechanics' physics optimization, and MRI-personalization. Compared to the main text, we show these results separately per gender. The averages differ slightly between genders. In Table A.6, we show the residual forces and moments for all models, separated by gender. Here, female participants show lower residual forces and moments accross all models, however, modelling errors are not the only gender-dependent factor. When comparing the effect of personalization on metabolics per gender, we found only slight differences between the two groups.

Table A.5: **Results of the MRIgait benchmark for participants by gender.** The first two row show state-of-the-art methods of scaling (scaled) and addBiomechanics' physics optimization (addB). The next two rows show the novel SIPP-personalized and scaled SIPP-generic (S-gen) models. In the final two rows, SIPP and SIPP-generic in combination with addBiomechanics' physics optimization are shown (add+SIPP, add+S-gen). The columns show the mean and standard deviation of the mean absolute deviation (MAD) and mean relative deviation (MRD) for mass (M), center of mass (CoM), inertia (I), and kinetic energy (E). The best results are highlighted in bold. Inertia values only account for the diagonal elements of the inertia tensor, as generic OpenSim models assume off-diagonal elements to be zero.

| | **Mean Absolute Deviation** | | | | **Mean Relative Deviation** | | | |
|---|---|---|---|---|---|---|---|---|
| **Model** | **M** [g] | **CoM** [cm] | **I** [kg m$^2$] | **E** [J] | **M** [%] | **CoM** [%] | **I** [%] | **E** [%] |
| | | | | **Male Participants** | | | | |
| scaled | $80.7 \pm 92.2$ | $9.5 \pm 0.8$ | $0.38 \pm 0.60$ | $2.3 \pm 0.7$ | $22.3 \pm 6.7$ | $56.7 \pm 2.0$ | $28.1 \pm 13.2$ | $22.2 \pm 9.1$ |
| SIPP | $56.4 \pm 71.6$ | $4.9 \pm 1.6$ | $0.34 \pm 0.58$ | $1.8 \pm 0.6$ | $14.7 \pm 2.7$ | $23.6 \pm 4.5$ | $22.1 \pm 1.8$ | $16.9 \pm 6.3$ |
| S-gen | $35.8 \pm 261.5$ | $5.0 \pm 1.7$ | $0.35 \pm 0.56$ | $1.5 \pm 0.5$ | $6.5 \pm 3.7$ | $23.5 \pm 5.3$ | $4.9 \pm 3.5$ | $13.7 \pm 6.0$ |
| addB | $45.3 \pm 144.8$ | $9.5 \pm 0.6$ | $0.40 \pm 0.59$ | $1.7 \pm 0.5$ | $15.7 \pm 9.8$ | $56.8 \pm 2.5$ | $25.1 \pm 9.9$ | $16.9 \pm 7.8$ |
| addB+SIPP | $44.4 \pm 80.9$ | $4.9 \pm 1.7$ | $0.36 \pm 0.59$ | $2.3 \pm 0.5$ | $22.1 \pm 0.6$ | $23.2 \pm 5.3$ | $26.7 \pm 7.0$ | $21.5 \pm 4.0$ |
| addB+S-gen | $76.2 \pm 323.6$ | $5.7 \pm 1.7$ | $0.35 \pm 0.58$ | $1.1 \pm 0.4$ | $7.6 \pm 3.7$ | $27.2 \pm 4.5$ | $2.8 \pm 2.8$ | $9.8 \pm 3.7$ |
| | | | | **Female Participants** | | | | |
| scaled | $83.5 \pm 44.0$ | $7.9 \pm 0.5$ | $0.32 \pm 0.58$ | $2.0 \pm 0.5$ | $36.9 \pm 2.5$ | $65.1 \pm 1.9$ | $51.2 \pm 4.7$ | $32.3 \pm 4.5$ |
| SIPP | $56.4 \pm 35.4$ | $1.3 \pm 0.2$ | $0.38 \pm 0.13$ | $0.2 \pm 0.2$ | $4.6 \pm 3.5$ | $9.9 \pm 0.8$ | $8.7 \pm 3.4$ | $4.0 \pm 3.1$ |
| S-gen | $89.7 \pm 18.1$ | $1.9 \pm 0.2$ | $0.24 \pm 0.12$ | $0.4 \pm 0.4$ | $2.2 \pm 1.0$ | $12.8 \pm 1.6$ | $7.1 \pm 5.8$ | $4.9 \pm 4.5$ |
| addB | $20.5 \pm 13.2$ | $8.0 \pm 0.6$ | $0.37 \pm 0.70$ | $1.5 \pm 0.4$ | $30.9 \pm 4.6$ | $66.3 \pm 2.0$ | $43.8 \pm 7.9$ | $23.9 \pm 2.9$ |
| add+SIPP | $63.1 \pm 64.9$ | $3.9 \pm 4.1$ | $0.19 \pm 0.32$ | $1.1 \pm 0.5$ | $14.5 \pm 10.1$ | $31.1 \pm 31.7$ | $24.2 \pm 21.9$ | $17.1 \pm 6.6$ |
| add+S-gen | $107.7 \pm 63.6$ | $1.7 \pm 0.3$ | $0.15 \pm 0.55$ | $0.6 \pm 0.4$ | $2.6 \pm 1.2$ | $12.3 \pm 2.6$ | $3.3 \pm 1.5$ | $7.6 \pm 3.6$ |

#### Misgendered SIPP-generic

In the main text, we presented the results of the SIPP-generic model. To underline the importance of gender-specific BSIP parameterizations, we show the results of the SIPP-generic model when misgendered in Table A.7. The results show that the SIPP-generic model performs slightly worse than the scaled model when misgendered.

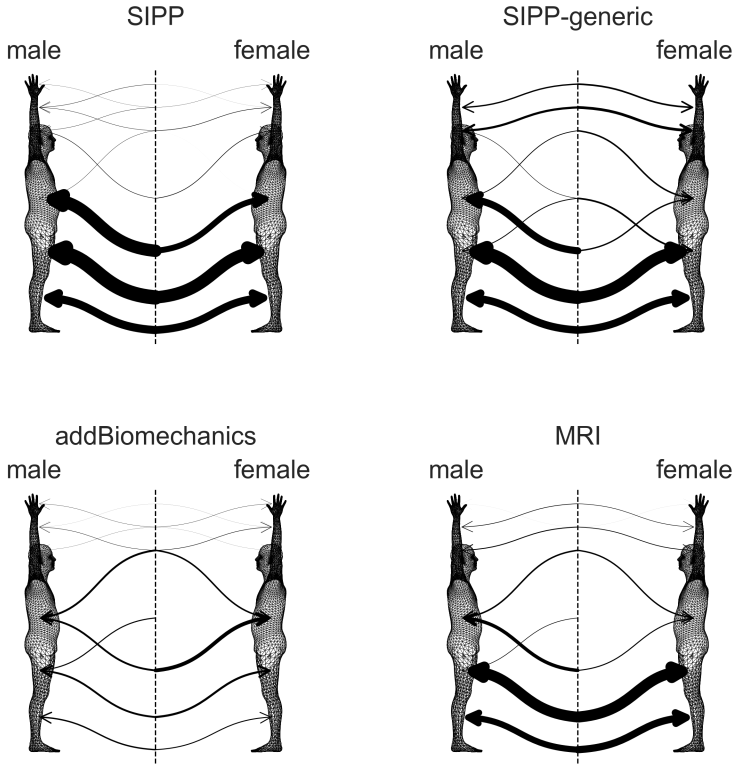


Figure A.9: **Average mass reallocation compared to scaled models.** The arrows depict the average transfer of mass from the scaled (dotted line in the middle) to the personalized (male: left, female: right) MSK models. We binned the body parts into 7 regions, which are (from top to bottom): hand, lower arm, upper arm, trunk, upper leg, lower leg and feet. The thickness of the arrows corresponds to the average amount of mass reallocated from one segment to another.

Table A.6: **Overview of residual forces.** We show the mean and standard deviation of the absolute residuals for all dimensions ($x$, $y$, $z$) and their norms. Forces are listed as a percentage of bodyweight [BW%] and the values for absolute residual moments are listed as a percentage of bodyweight times bodyweight [BWBH%]. We show the results for the MRI-personalized model, scaled model, SIPP model, and SIPP-generic (S-gen) model, in the first four rows. In the last three rows, we show the results for the same models in combination with addBiomechanics' physics optimization (addB, add+SIPP, add+S-gen). The best results per table section are highlighted in bold.

| | **Forces** | | | | **Moments** | | | |
|---|---|---|---|---|---|---|---|---|
| **Model** | $\mathbf{F}_x$ [BW%] | $\mathbf{F}_y$ [BW%] | $\mathbf{F}_z$ [BW%] | $\lVert\mathbf{F}\rVert$ [BW%] | $\mathbf{M}_x$ [BWBH%] | $\mathbf{M}_y$ [BWBH%] | $\mathbf{M}_z$ [BWBH%] | $\lVert\mathbf{M}\rVert$ [BWBH%] |
| **Male Participants** | | | | | | | | |
| MRI | $6.51 \pm 10.1$ | $3.95 \pm 4.6$ | $3.41 \pm 3.5$ | $9.58 \pm 10.6$ | $2.53 \pm 2.8$ | $1.04 \pm 1.0$ | $2.38 \pm 8.3$ | $4.09 \pm 8.6$ |
| scaled | $6.25 \pm 6.6$ | $4.05 \pm 3.8$ | $2.98 \pm 2.9$ | $9.05 \pm 6.9$ | $2.66 \pm 2.5$ | $1.10 \pm 0.9$ | $\mathbf{1.56 \pm 7.3}$ | $3.67 \pm 7.6$ |
| SIPP | $\mathbf{4.97 \pm 6.5}$ | $\mathbf{3.33 \pm 3.5}$ | $\mathbf{2.95 \pm 2.8}$ | $\mathbf{7.67 \pm 6.9}$ | $\mathbf{1.89 \pm 1.7}$ | $\mathbf{0.96 \pm 0.8}$ | $1.62 \pm 7.3$ | $\mathbf{3.02 \pm 7.4}$ |
| S-gen | $6.21 \pm 9.6$ | $3.94 \pm 4.6$ | $3.46 \pm 3.6$ | $9.40 \pm 10.2$ | $2.26 \pm 2.5$ | $1.01 \pm 1.0$ | $2.04 \pm 8.2$ | $3.66 \pm 8.4$ |
| addB | $3.80 \pm 7.0$ | $3.57 \pm 4.5$ | $\mathbf{1.61 \pm 3.1}$ | $6.12 \pm 8.4$ | $2.20 \pm 1.8$ | $1.08 \pm 1.0$ | $1.47 \pm 8.2$ | $3.22 \pm 8.3$ |
| addB+SIPP | $\mathbf{2.95 \pm 5.6}$ | $\mathbf{3.01 \pm 4.5}$ | $1.70 \pm 4.1$ | $\mathbf{5.20 \pm 7.9}$ | $\mathbf{1.54 \pm 1.4}$ | $\mathbf{0.95 \pm 1.0}$ | $\mathbf{1.45 \pm 8.1}$ | $\mathbf{2.62 \pm 8.2}$ |
| addB+S-gen | $3.08 \pm 6.2$ | $3.18 \pm 4.6$ | $1.68 \pm 4.6$ | $5.40 \pm 8.6$ | $1.75 \pm 1.4$ | $0.99 \pm 1.0$ | $1.70 \pm 8.2$ | $2.96 \pm 8.2$ |
| **Female Participants** | | | | | | | | |
| MRI | $3.23 \pm 3.8$ | $3.21 \pm 3.7$ | $2.75 \pm 3.0$ | $6.14 \pm 5.2$ | $1.08 \pm 1.0$ | $0.79 \pm 0.6$ | $1.50 \pm 1.5$ | $2.28 \pm 1.6$ |
| scaled | $4.25 \pm 3.9$ | $3.24 \pm 3.1$ | $\mathbf{2.42 \pm 2.6}$ | $6.66 \pm 4.6$ | $1.52 \pm 1.3$ | $0.82 \pm 0.7$ | $\mathbf{1.04 \pm 1.3}$ | $2.27 \pm 1.6$ |
| SIPP | $\mathbf{3.00 \pm 3.3}$ | $\mathbf{2.94 \pm 3.2}$ | $2.58 \pm 2.7$ | $\mathbf{5.68 \pm 4.5}$ | $\mathbf{1.03 \pm 1.0}$ | $\mathbf{0.76 \pm 0.6}$ | $1.32 \pm 1.3$ | $\mathbf{2.09 \pm 1.4}$ |
| S-gen | $3.30 \pm 3.8$ | $3.34 \pm 3.8$ | $2.99 \pm 3.2$ | $6.42 \pm 5.4$ | $1.10 \pm 1.0$ | $0.82 \pm 0.7$ | $1.60 \pm 1.5$ | $2.38 \pm 1.6$ |
| addB | $3.42 \pm 4.8$ | $3.37 \pm 4.6$ | $\mathbf{1.72 \pm 2.7}$ | $5.73 \pm 6.7$ | $1.56 \pm 1.4$ | $0.78 \pm 0.7$ | $\mathbf{1.02 \pm 1.3}$ | $2.28 \pm 1.7$ |
| addB+SIPP | $\mathbf{2.56 \pm 4.4}$ | $\mathbf{3.15 \pm 4.2}$ | $1.87 \pm 2.7$ | $\mathbf{5.10 \pm 6.2}$ | $1.18 \pm 1.3$ | $\mathbf{0.78 \pm 0.6}$ | $1.23 \pm 1.4$ | $\mathbf{2.15 \pm 1.7}$ |
| addB+S-gen | $2.60 \pm 4.0$ | $3.31 \pm 4.5$ | $1.93 \pm 3.0$ | $5.29 \pm 6.2$ | $\mathbf{1.07 \pm 1.0}$ | $0.78 \pm 0.6$ | $1.50 \pm 1.5$ | $2.25 \pm 1.6$ |

Table A.7: **Overview of residual forces of SIPP-generic and SIPP-generic when the wrong gender is assigned.** We show the mean and standard deviation of the absolute residuals for all dimensions ($x$, $y$, $z$) and their norms. Forces are listed as a percentage of bodyweight [BW%] and the values for absolute residual moments are listed as a percentage of bodyweight times bodyweight [BWBH%]. The best results are highlighted in bold.

| | **Forces** | | | | **Moments** | | | |
|---|---|---|---|---|---|---|---|---|
| **Model** | $\mathbf{F}_x$ [BW%] | $\mathbf{F}_y$ [BW%] | $\mathbf{F}_z$ [BW%] | $\lVert\mathbf{F}\rVert$ [BW%] | $\mathbf{M}_x$ [BWBH%] | $\mathbf{M}_y$ [BWBH%] | $\mathbf{M}_z$ [BWBH%] | $\lVert\mathbf{M}\rVert$ [BWBH%] |
| S-gen | $\mathbf{4.66 \pm 6.8}$ | $\mathbf{3.56 \pm 4.0}$ | $\mathbf{3.19 \pm 3.4}$ | $\mathbf{7.77 \pm 7.6}$ | $1.64 \pm 1.7$ | $0.92 \pm 0.8$ | $\mathbf{1.83 \pm 4.8}$ | $\mathbf{3.00 \pm 4.9}$ |
| S-gen, misgendered | $4.74 \pm 7.2$ | $3.57 \pm 4.1$ | $3.24 \pm 3.3$ | $7.88 \pm 8.0$ | $\mathbf{1.64 \pm 1.8}$ | $\mathbf{0.91 \pm 0.8}$ | $1.84 \pm 4.8$ | $3.02 \pm 5.0$ |
